# A comparative analysis of chromatin accessibility in cattle, pig, and mouse tissues

**DOI:** 10.1101/2020.08.13.249870

**Authors:** Michelle M Halstead, Colin Kern, Perot Saelao, Ying Wang, Ganrea Chanthavixay, Juan F Medrano, Alison L Van Eenennaam, Ian Korf, Christopher K Tuggle, Catherine W Ernst, Huaijun Zhou, Pablo J Ross

## Abstract

**Background:** Although considerable progress has been made towards annotating the noncoding portion of the human and mouse genomes, regulatory elements in other species, such as livestock, remain poorly characterized. This lack of functional annotation poses a substantial roadblock to agricultural research and diminishes the value of these species as model organisms. As active regulatory elements are typically characterized by chromatin accessibility, we implemented the Assay for Transposase Accessible Chromatin (ATAC-seq) to annotate and characterize regulatory elements in pigs and cattle, given a set of eight adult tissues.

**Results:** Overall, 306,304 and 273,594 active regulatory elements were identified in pig and cattle, respectively. 71,478 porcine and 47,454 bovine regulatory elements were highly tissue-specific and were correspondingly enriched for binding motifs of known tissue-specific transcription factors. However, in every tissue the most prevalent accessible motif corresponded to the insulator CTCF, suggesting pervasive involvement in 3-D chromatin organization. Taking advantage of a similar dataset in mouse, open chromatin in pig, cattle, and mice were compared, revealing that the conservation of regulatory elements, in terms of sequence identity and accessibility, was consistent with evolutionary distance; whereas pig and cattle shared about 20% of accessible sites, mice and ungulates only had about 10% of accessible sites in common. Furthermore, conservation of accessibility was more prevalent at promoters than at intergenic regions.

**Conclusions:** The lack of conserved accessibility at distal elements is consistent with rapid evolution of enhancers, and further emphasizes the need to annotate regulatory elements in individual species, rather than inferring elements based on homology. This atlas of chromatin accessibility in cattle and pig constitutes a substantial step towards annotating livestock genomes and dissecting the regulatory link between genome and phenome.

## Background

Despite considerable progress to annotate protein-coding genes in livestock species, the vast majority of these genomes is noncoding and remains poorly characterized. Epigenomics techniques, such as chromatin immunoprecipitation followed by sequencing (ChIP-seq) and DNase I hypersensitive sites sequencing (DNase-seq), have been extensively employed to catalog functional elements in humans (1) and classical model organisms (2–6). For instance, the international human epigenome consortium has profiled thousands of epigenomes and identified millions of regulatory elements in the human genome (1,7,8), yielding an atlas of functional elements that has been invaluable for subsequent research in a wide variety of biological processes, including disease (9–13), pluripotency (14–16), differentiation (17–19), and morphology (20,21).

Ultimately, genome-wide patterns of chromatin accessibility and compaction determine which genomic regions are available to cellular machinery, and are thereby intimately connected to the cell-specific gene expression patterns that determine identity and function. Controlled exposure of specific sites provides opportunities for transcription factors to bind their recognition motifs and regulate gene expression through further recruitment of proteins, such as RNA polymerases (22,23). Consequently, profiling open chromatin has the potential to not only identify regulatory elements, but also profile their activities in different cell types. The increasing availability of next-generation sequencing-based techniques spurred development of several alternative epigenomics assays, such as the Assay for Transposase Accessible Chromatin (ATAC-seq), which was first reported by Buenrostro *et al* in 2013 (24). Following its introduction, ATAC-seq quickly became one of the leading methods for identification of open chromatin, largely due to the simplicity of the technique and low input requirements, which made it possible to study chromatin structure in rare samples.

Here we implemented ATAC-seq to profile open chromatin in a set of cattle and pig tissues: subcutaneous adipose, brain (frontal brain cortex, hypothalamus, and cerebellum), liver, lung, skeletal muscle, and spleen. This set of prioritized tissues are associated with a large number of qualitative phenotypic traits relevant to animal production, such as disease resistance, growth, and feed efficiency. Overall, about 300,000 accessible regions were identified in each species, yielding an epigenomic resource that will benefit agricultural genomics research and enable cross-species comparisons that will enhance knowledge of comparative epigenomics and transcriptional regulation.

## Results

### ATAC-seq library quality control and preprocessing

Using a modified ATAC-seq protocol, genome-wide chromatin accessibility was profiled in eight tissues derived from two adult male Hereford cattle and two adult male Yorkshire pigs: three brain tissues (frontal brain cortex, hypothalamus, and cerebellum), liver, lung, spleen, subcutaneous adipose, and skeletal muscle (Figure 1a). With the exception of one cattle cerebellum sample, which was lost during processing, ATAC-seq data were generated for two biological replicates per tissue (Table S1). In addition, two technical replicates were prepared for cattle cortex, pig cerebellum, and pig hypothalamus. ATAC-seq signal from technical replicates were highly correlated (Pearson R averaged 0.97), and principal components analysis (PCA) of genome-wide signal grouped biological and technical replicates together (Figure S1).

**Figure 1.**
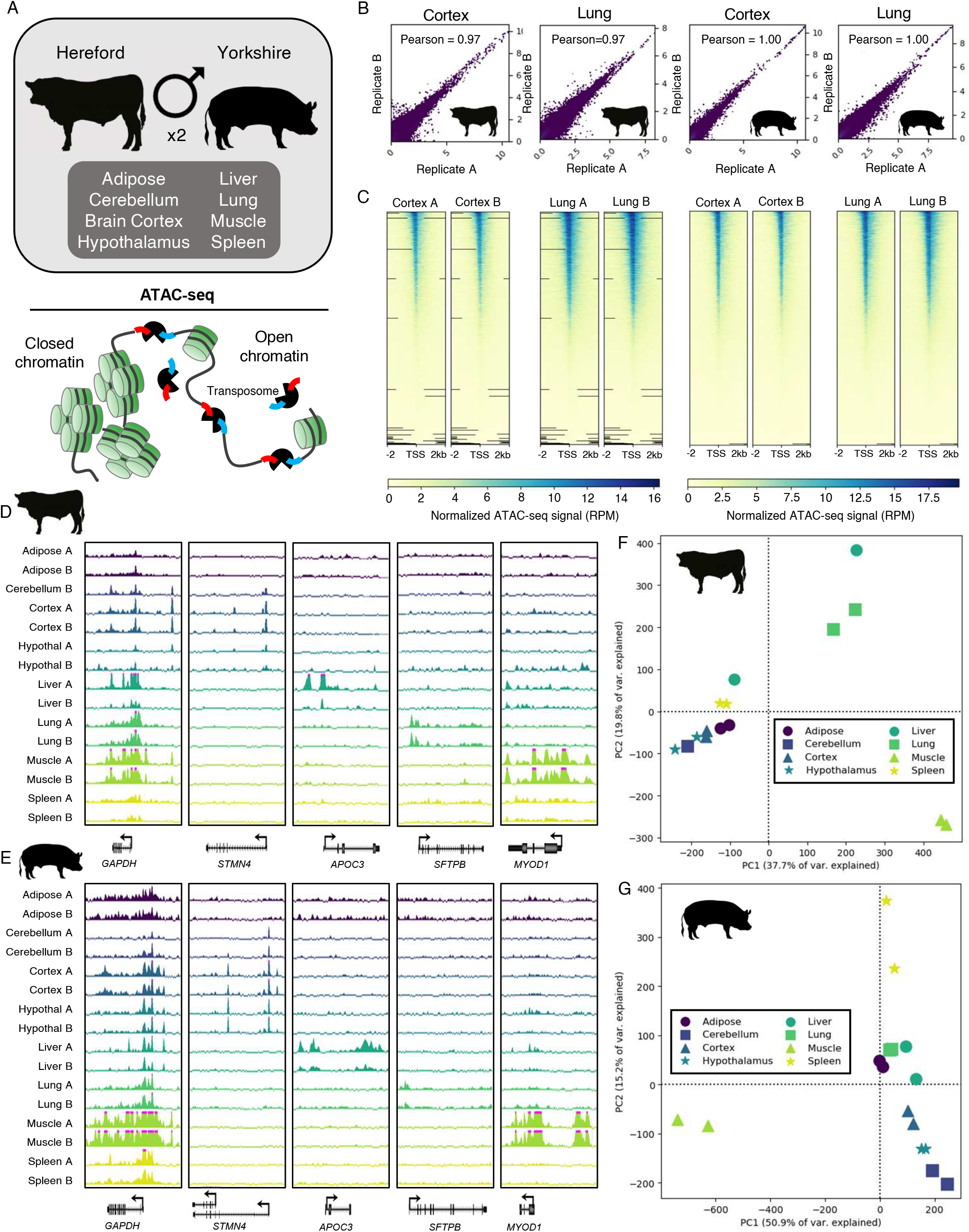
Experimental design and ATAC-seq quality metrics. A) Overview of tissues collected from adult male cattle and pigs for ATAC-seq, wherein the Tn5 transposase preferentially cuts DNA at accessible sites and simultaneously inserts sequencing adapters. B) Scatterplots showing Pearson correlation of normalized genome-wide ATAC-seq signal (reads per million; RPM) between biological replicates for brain cortex and lung. C) Heatmaps depicting normalized ATAC-seq signal at all TSS, sorted by signal intensity. Signal shown for brain cortex and lung. D) Normalized ATAC-seq signal in cattle tissues at the GAPDH locus (a housekeeping gene), as well as several genes with tissue-specific activity. Pink bars indicate that signal exceeded the viewing range (up to 20 RPM). E) Normalized ATAC-seq signal in pig tissues at the same genes. F) PCA of normalized ATAC-seq signal in consensus open chromatin identified in cattle. G) PCA of normalized ATAC-seq signal in consensus open chromatin identified in pig.

Mapping of sequencing data resulted in 58 ± 4 million informative reads (uniquely mapping, non-mitochondrial, monoclonal reads) per sample (Table 1). Normalized genome-wide ATAC-seq signal was highly reproducible between biological replicates, with the Pearson correlation coefficient averaging 0.97 (Figure 1b; Figure S2), and increased in intensity at transcription start sites (TSS) (Figure 1c; Figure S3). Genes with tissue-specific functions demonstrated open chromatin that was specific to the corresponding tissue; for instance, the gene STMN4, which is involved in neuron projection development, is specifically marked by open chromatin at two sites in both cattle (Figure 1d) and pig (Figure 1e) in all three brain tissues, which likely correspond to alternate TSS, based on the gene annotation in pig. PCA of normalized ATAC-seq signal separated samples by tissue, with brain tissues grouping together in both pig and cattle (Figure 1f,g, Figure S4).

**Table 1.** ATAC-seq data preprocessing. Per library, total raw reads, mapped reads (excluding mitochondrial DNA), duplicate reads, informative reads (monoclonal and uniquely mapping), and percent of raw reads that were informative.

| *Species* | *Tissue* | *Rep* | *Raw reads* | *Mapped reads* | *Duplicate reads* | *Informative reads* | *% Raw* |
| --- | --- | --- | --- | --- | --- | --- | --- |
| Cattle | Adipose | A | 152,184,070 | 145,648,161 | 82,972,324 | 37,815,392 | 24.85 |
|  | Adipose | B | 126,920,776 | 123,879,749 | 25,466,938 | 72,623,985 | 57.22 |
|  | Cerebellum | B | 158,964,554 | 134,391,900 | 53,303,886 | 49,305,250 | 31.02 |
|  | Brain Cortex | A | 216,202,716 | 189,763,599 | 122,567,574 | 30,997,045 | 14.34 |
|  | Brain Cortex | B | 205,363,760 | 174,961,995 | 96,411,128 | 41,021,487 | 19.98 |
|  | Hypothalamus | A | 146,439,048 | 102,771,471 | 50,044,176 | 35,090,140 | 23.96 |
|  | Hypothalamus | B | 68,153,356 | 63,771,723 | 31,495,325 | 15,328,535 | 22.49 |
|  | Liver | A | 175,124,072 | 165,112,503 | 95,001,248 | 40,031,560 | 22.86 |
|  | Liver | B | 194,367,992 | 179,265,341 | 99,942,567 | 36,472,289 | 18.76 |
|  | Lung | A | 163,469,102 | 159,710,295 | 27,682,063 | 99,691,291 | 60.98 |
|  | Lung | B | 190,030,170 | 184,571,092 | 36,657,121 | 110,584,502 | 58.19 |
|  | Muscle | A | 89,174,618 | 86,045,857 | 15,262,767 | 55,794,693 | 62.57 |
|  | Muscle | B | 97,247,280 | 69,592,640 | 10,539,048 | 45,909,945 | 47.21 |
|  | Spleen | A | 261,785,316 | 250,911,567 | 163,860,741 | 37,521,241 | 14.33 |
|  | Spleen | B | 179,854,284 | 162,449,025 | 51,533,284 | 69,626,100 | 38.71 |
| Pig | Adipose | A | 93,118,998 | 87,025,520 | 15,814,107 | 57,651,677 | 61.91 |
|  | Adipose | B | 71,639,956 | 66,597,412 | 12,368,404 | 42,960,583 | 59.97 |
|  | Cerebellum | A | 186,388,542 | 175,280,070 | 90,013,906 | 63,538,783 | 34.09 |
|  | Cerebellum | B | 125,817,350 | 116,536,743 | 41,557,703 | 57,997,395 | 46.10 |
|  | Brain Cortex | A | 101,924,240 | 98,384,411 | 36,110,627 | 53,096,952 | 52.09 |
|  | Brain Cortex | B | 160,871,726 | 155,783,257 | 77,377,043 | 66,403,810 | 41.28 |
|  | Hypothalamus | A | 112,463,726 | 106,966,835 | 52,803,730 | 39,919,895 | 35.50 |
|  | Hypothalamus | B | 170,163,006 | 162,907,052 | 92,303,710 | 56,125,160 | 32.98 |
|  | Liver | A | 171,864,386 | 167,321,556 | 113,588,699 | 42,376,220 | 24.66 |
|  | Liver | B | 169,952,062 | 164,205,920 | 85,279,200 | 64,117,458 | 37.73 |
|  | Lung | A | 108,086,464 | 104,424,556 | 19,658,156 | 73,018,749 | 67.56 |
|  | Lung | B | 110,690,180 | 106,275,981 | 17,791,030 | 74,869,852 | 67.64 |
|  | Muscle | A | 168,211,474 | 165,225,305 | 41,747,860 | 113,058,649 | 67.21 |
|  | Muscle | B | 141,069,686 | 138,785,466 | 39,577,315 | 91,780,486 | 65.06 |
|  | Spleen | A | 91,549,652 | 88,698,199 | 13,254,769 | 64,352,893 | 70.29 |
|  | Spleen | B | 93,747,292 | 90,558,895 | 12,817,045 | 67,241,934 | 71.73 |

Several commonly used statistics were used to evaluate library quality (Table 2). The non-redundant read fraction (NRF), which gauges library complexity by measuring the proportion of non-duplicate uniquely mapped reads out of all mapped reads, averaged 0.62 ± 0.03, indicating acceptable library complexity. The synthetic Jensen-Shannon distance (sJSD), which measures the divergence between the genome-wide ATAC-seq signal in a given sample versus a uniform distribution, averaged 0.46 ± 0.01, suggesting a non-random ATAC-seq signal distribution throughout the genome. Finally, the Fraction of Reads in Peaks (FRiP) was calculated to evaluate the strength of signal over background. On average, 130,712 ± 10,994 ATAC-seq peaks (regions of enrichment) were called per sample (Supplementary Data 1-2), and the fraction of reads in peaks (FRiP score) averaged 34.8 ± 2.6%. Notably, FRiP scores for adipose libraries were consistently lower (average 8.5 ± 2.5%) than for other tissues (average 38.7 ± 2.1%).

**Table 2.** Quality metrics of ATAC-seq libraries. Non-redundant read fraction (NRF) measures library complexity, Fraction of reads in peaks (FRiP) measures signal-to-noise ratio, and synthetic Jensen-Shannon distance (sJSD) measures divergence between ATAC-seq signal and a uniform distribution.

| *Species* | *Tissue* | *Rep.* | *NRF* | *FRiP* | *sJSD* |
| --- | --- | --- | --- | --- | --- |
| Cattle | Adipose | A | 0.43 | 9.82 | 0.34 |
|  | Adipose | B | 0.79 | 15.04 | 0.34 |
|  | Cerebellum | B | 0.60 | 42.53 | 0.48 |
|  | Brain Cortex | A | 0.35 | 30.16 | 0.48 |
|  | Brain Cortex | B | 0.45 | 40.80 | 0.49 |
|  | Hypothalamus | A | 0.51 | 43.67 | 0.50 |
|  | Hypothalamus | B | 0.51 | 11.97 | 0.37 |
|  | Liver | A | 0.42 | 37.99 | 0.50 |
|  | Liver | B | 0.44 | 28.17 | 0.44 |
|  | Lung | A | 0.83 | 37.63 | 0.45 |
|  | Lung | B | 0.80 | 44.26 | 0.48 |
|  | Muscle | A | 0.82 | 40.77 | 0.50 |
|  | Muscle | B | 0.85 | 41.77 | 0.51 |
|  | Spleen | A | 0.35 | 21.29 | 0.42 |
|  | Spleen | B | 0.68 | 35.24 | 0.45 |
| Pig | Adipose | A | 0.82 | 5.21 | 0.40 |
|  | Adipose | B | 0.81 | 4.12 | 0.40 |
|  | Cerebellum | A | 0.49 | 43.05 | 0.46 |
|  | Cerebellum | B | 0.64 | 41.49 | 0.47 |
|  | Brain Cortex | A | 0.63 | 40.65 | 0.48 |
|  | Brain Cortex | B | 0.50 | 44.53 | 0.46 |
|  | Hypothalamus | A | 0.51 | 36.85 | 0.49 |
|  | Hypothalamus | B | 0.43 | 37.76 | 0.44 |
|  | Liver | A | 0.32 | 55.38 | 0.52 |
|  | Liver | B | 0.48 | 49.25 | 0.49 |
|  | Lung | A | 0.81 | 31.21 | 0.43 |
|  | Lung | B | 0.83 | 36.87 | 0.46 |
|  | Muscle | A | 0.75 | 58.16 | 0.62 |
|  | Muscle | B | 0.71 | 61.94 | 0.65 |
|  | Spleen | A | 0.85 | 29.41 | 0.43 |
|  | Spleen | B | 0.86 | 22.70 | 0.38 |

For each tissue, peaks called from biological replicates were compared to evaluate consistency between replicates and identify accessible regions with high confidence. On average, 67.4 ± 17.8% of peaks from the replicate with fewer peaks called were also identified in the other replicate (Table 3). Regions that were enriched for ATAC-seq signal in both biological replicates of at least one tissue were termed ‘consensus’ peaks. In the case of cattle cerebellum, for which only one biological replicate was available, ATAC-seq peaks were called more stringently to identify consensus peaks. Altogether, 306,304 and 273,594 consensus peaks were identified in pig and cattle, respectively (Table 4).

**Table 3.** Replicability of ATAC-seq peaks.

| *Species* | *Tissue* | *Rep A Peaks* | *Rep B Peaks* | *Rep A Peaks overlapping Rep B Peaks* | *% Rep A Peaks* | *Rep B Peaks overlapping Rep A peaks* | *% Rep B Peaks* |
| --- | --- | --- | --- | --- | --- | --- | --- |
| Cattle | Adipose | 59,612 | 133,768 | 38,647 | 64.8 | 38,471 | 28.8 |
|  | Cortex | 109,395 | 160,546 | 75,462 | 69.0 | 75,206 | 46.8 |
|  | Hypothalamus | 59,966 | 37,036 | 19,970 | 33.3 | 19,999 | 54.0 |
|  | Liver | 102,704 | 114,583 | 58,444 | 56.9 | 58,563 | 51.1 |
|  | Lung | 221,576 | 248,844 | 167,311 | 75.5 | 166,777 | 67.0 |
|  | Muscle | 107,208 | 113,502 | 76,780 | 71.6 | 76,801 | 67.7 |
|  | Spleen | 110,852 | 200,323 | 79,355 | 71.6 | 79,007 | 39.4 |
| Pig | Adipose | 9,192 | 7,645 | 4,778 | 52.0 | 4,778 | 62.5 |
|  | Cerebellum | 220,327 | 214,455 | 132,123 | 60.0 | 132,292 | 61.7 |
|  | Cortex | 142,658 | 156,069 | 103,581 | 72.6 | 103,382 | 66.2 |
|  | Hypothalamus | 103,402 | 144,418 | 63,382 | 61.3 | 63,368 | 43.9 |
|  | Liver | 112,202 | 136,085 | 78,395 | 69.9 | 78,274 | 57.5 |
|  | Lung | 167,298 | 191,926 | 114,470 | 68.4 | 113,864 | 59.3 |
|  | Muscle | 137,000 | 105,823 | 92,039 | 67.2 | 92,705 | 87.6 |
|  | Spleen | 100,982 | 100,867 | 64,192 | 63.6 | 64,208 | 63.7 |

**Table 4.** Consensus ATAC-seq peaks identified for each tissue in pig, cattle, and mouse. To obtain a single comprehensive set of unique intervals of open chromatin, accounting for accessibility detected in any of the eight tissues, consensus peaks from each tissue were combined such that any overlapping peaks were merged into a single peak.

| *Tissue* | *Pig* | *Cattle* | *Mouse* |
| --- | --- | --- | --- |
| Adipose | 4,785 | 38,745 | 64,008 |
| Cerebellum | 134,086 | 93,927 | 57,771 |
| Cortex | 104,272 | 75,762 | 149,223 |
| Hypothalamus | 63,736 | 20,045 | -- |
| Liver | 78,841 | 58,853 | 90,228 |
| Lung | 115,491 | 169,734 | 76,168 |
| Muscle | 93,550 | 77,378 | 22,003 |
| Spleen | 64,667 | 79,960 | 35,278 |
| *Merged peak set* | 306,304 | 273,594 | 254,076 |

### Global characteristics of accessible chromatin in cattle, pig, and mouse

To infer the functional significance of accessible regions that were identified in pig and cattle tissues, consensus peaks were characterized by genomic localization (positioning relative to genes), sequence content, and tissue-specificity. In cattle, consensus peaks averaged 616 bp in width, and covered 6.2% of the genome. Similarly, pig consensus peaks were 624 bp wide, and accounted for 7.2% of the genome. As expected, open chromatin was particularly enriched near TSS compared to random sequences, although most consensus peaks were intronic or intergenic (Figure 2a). Consensus peaks were also frequently in only one tissue (54 and 58% of cattle and pig peaks, respectively) (Figure 2b).

**Figure 2.**
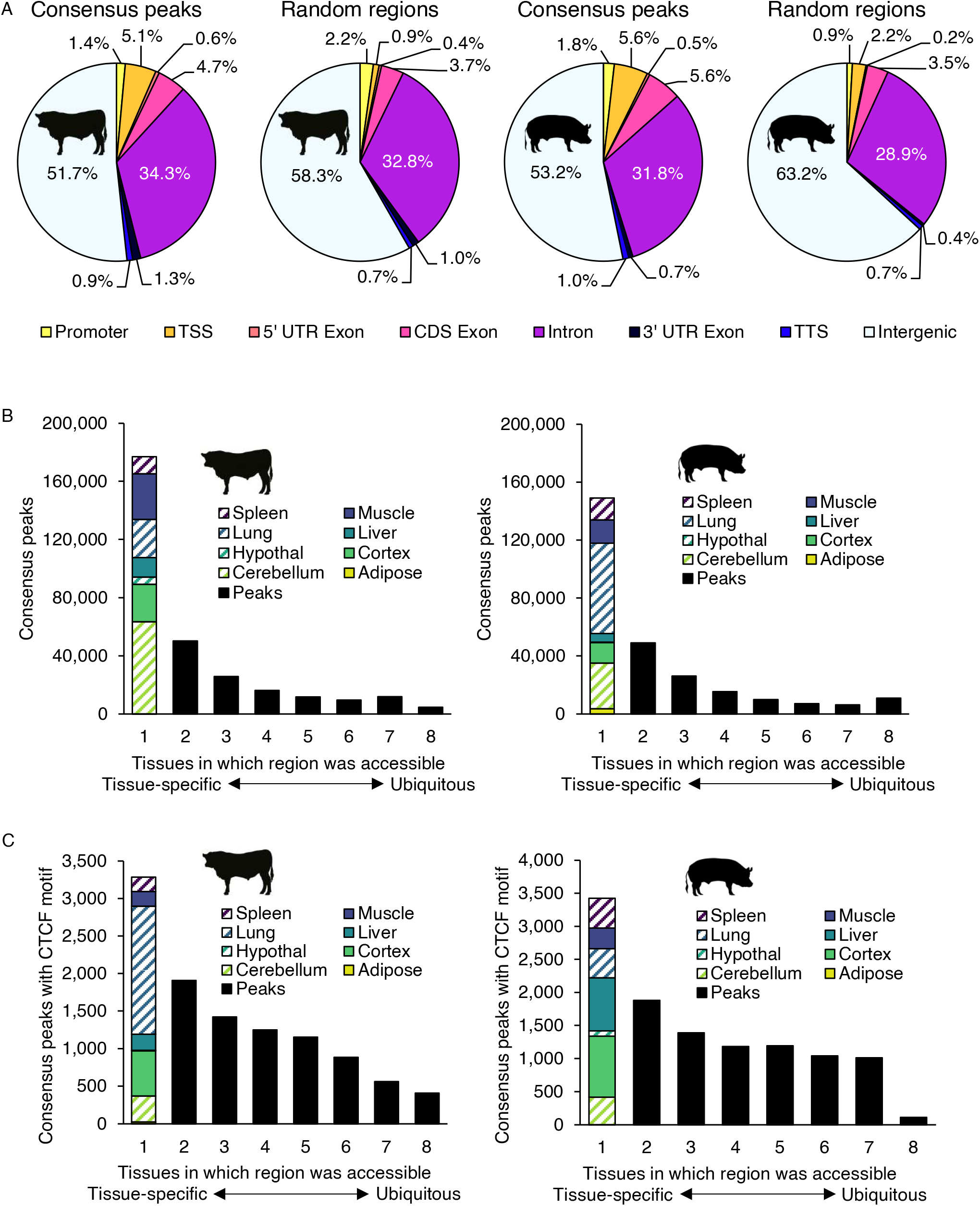
Open chromatin localization and differential accessibility. A) Distribution of cattle and pig consensus open chromatin, as well as randomized genomic intervals. Because peaks often span multiple features, peaks were categorized based on 1 bp overlap with features in the following order: first as TSS (±50 bp), then as promoter (2kb upstream of TSS), transcription termination site (TTS) (±50 bp), 5’ untranslated region (UTR), 3’ UTR, coding sequence (CDS), intronic, and if no features were overlapped, peaks were considered intergenic. B) Distribution of consensus peak activity, ranging from tissue-specific (accessible in only one tissue) to ubiquitous (accessible in all sampled tissues). Consensus peaks that were accessible in a single tissue were further broken down by tissue. C) Distribution of consensus peak activity for regions containing CTCF motifs.

A comparable mouse ATAC-seq dataset, which included libraries from two male replicates for all tissues except hypothalamus, was downloaded from the CNGB Nucleotide Sequence Read Archive (Project ID CNP0000198) and processed in the same manner as the pig and cattle data. Similar to cattle and pig, 254,076 consensus peaks were identified from mouse tissues (Table 4). Mouse consensus peaks also covered a comparable portion of the genome (6.2%), were of similar width (average 668 bp), demonstrated enrichment at TSS (Figure S5a), and were often tissue-specific (63% of peaks) (Figure S5b). Surprisingly, a higher fraction of mouse consensus peaks localized to TSS (9.6%) than in cattle (5.1%) or pig (5.6%) (Figure 2a, Figure S5a). This discrepancy could be attributed either to the protocol, as pig and cattle ATAC-seq libraries were subjected to size-selection and mouse libraries were not, or suboptimal genome annotations, as the pig and cattle annotations are relatively incomplete in comparison to the mouse. Nevertheless, the global characteristics of open chromatin were consistent across these three species.

To interrogate the potential function of accessible regions in cattle and pig, consensus peaks were subjected to motif enrichment analysis. Overall, consensus peaks were most significantly enriched for CTCF recognition sites, with about 8% of accessible regions harboring CTCF motifs in each species (Table 5). Of note, CTCF motifs were more prevalent in consensus peaks identified in 3 or more tissues (8% of peaks in cattle and pig) than in consensus peaks identified in only 1 or 2 tissues (3% and 2% of peaks in cattle and pig, respectively). Nevertheless, in both species 30% of consensus peaks containing a CTCF motif were only accessible in a single tissue (Figure 2c), indicating that CTCF binding could play both tissue-specific and ubiquitous regulatory roles.

**Table 5.**
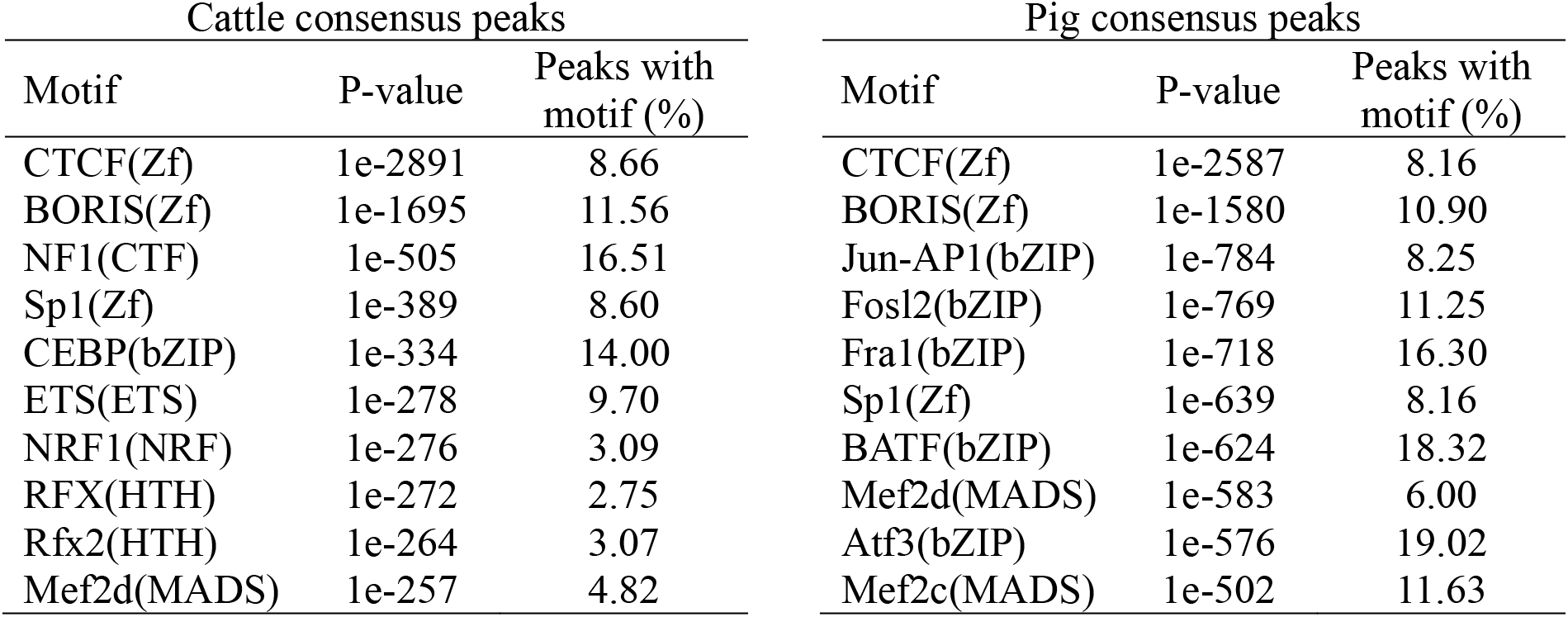
Motif enrichment in consensus open chromatin. Top ten enriched known binding motifs identified from the merged set of consensus peaks in each species.

The unique open chromatin landscapes present in different cell types are crucial for regulation of transcription, the products of which ultimately confer cell identity and function. Most consensus peaks (54% in cattle, 58% in pig, and 63% in mouse) were only present in a single tissue (Figure 2b), suggesting that these regions were involved in tissue-specific regulatory programs. These regions were of particular interest, considering that differentially accessible regions have been associated with higher density of transcription factor (TF) binding sites, hinting at interesting regulatory roles.

To stringently identify open chromatin that was specific to a given tissue, only consensus peaks that did not overlap any peaks called from either replicate in any other tissue were considered tissue-specific. In sum, 71,479 peaks in pig, 47,454 peaks in cattle, and 116,700 peaks in mouse (Figure 3a) were identified as having highly tissue-specific ATAC-seq signal (Figure 3b). Interestingly, tissues with the greatest number of tissue-specific peaks varied between species. Although cerebellum-specific peaks were numerous compared to the cortex in cattle and pig, the opposite trend was observed in the mouse. Of the remaining tissues, liver-specific peaks were particularly abundant in mouse, whereas lung-specific peaks were prevalent in cattle, and muscle-specific peaks were the most frequent in pig.

**Figure 3.**
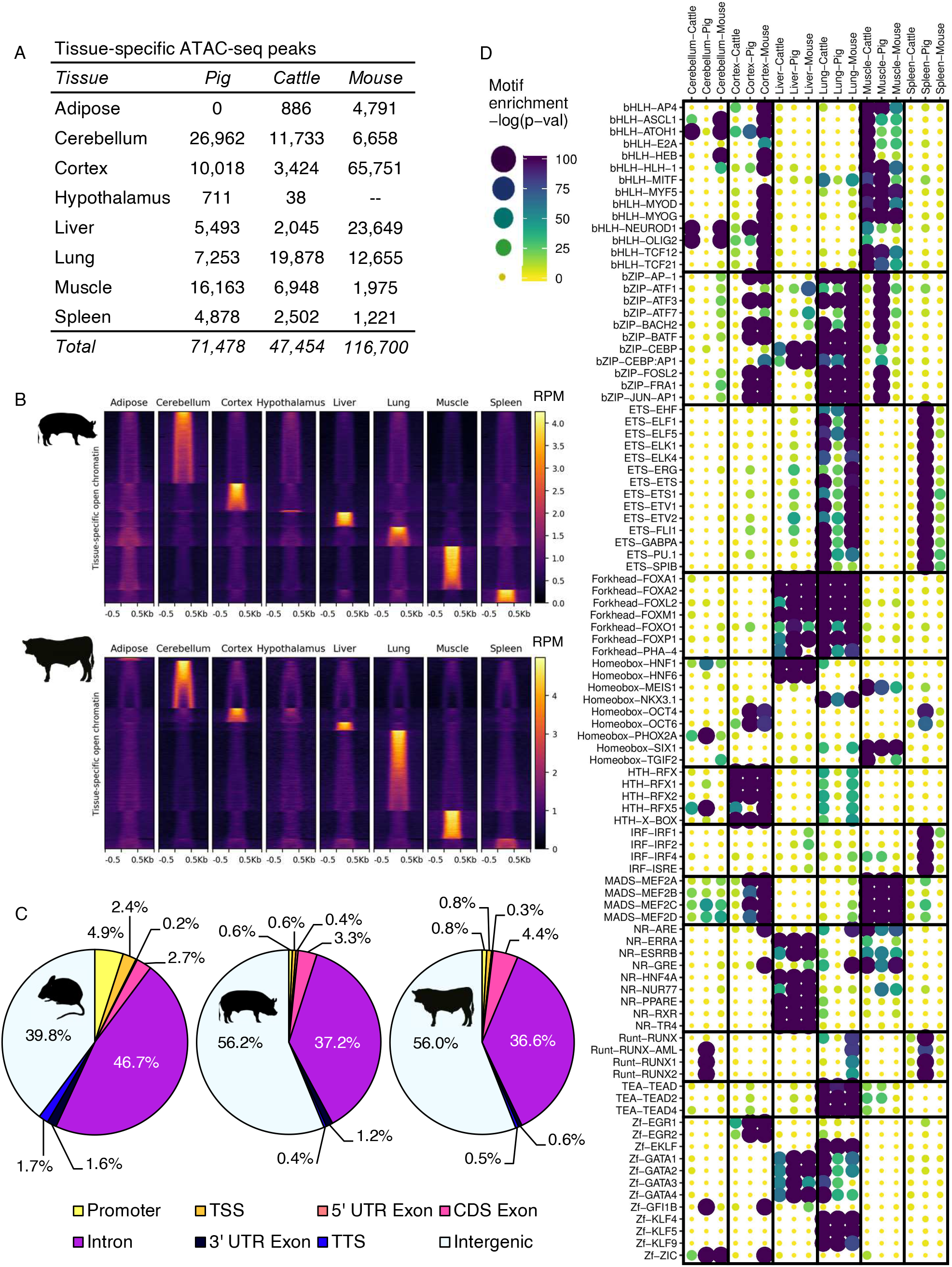
Characterization of highly tissue-specific open chromatin. A) Overview of regions demonstrating tissue-specific accessibility in each tissue and each species. B) Normalized ATAC-seq signal (RPM) at regions demonstrating tissue-specific accessibility in cattle and pig tissues. Tissue-specific peaks were first grouped by corresponding tissue, then ordered by signal intensity. Peaks were scaled to 500 bp, and signal is shown 500 bp upstream and downstream. C) Distribution of tissue-specific open chromatin relative to gene annotations. D) Enrichment of known TF binding motifs in tissue-specific open chromatin. Motifs sorted by TF family. Sets of tissue-specific open chromatin for each species are grouped by tissue. Increasing size and color intensity indicate increasing enrichment for a given motif.

Whereas all consensus peaks were enriched at TSS compared to random chance (Figure 2a, Figure S5a), tissue-specific peaks coincided with TSS less frequently than would be expected by random chance (Figure 3c). Although TSS annotation in these species is likely to be incomplete, the lack of tissue-specific peaks near annotated TSS suggests that tissue-specific open chromatin is more likely to delineate enhancers, which are known to demonstrate highly tissue-specific activity, both spatially and temporally. Tissue-specific peaks from cerebellum, cortex, liver, lung, muscle, and spleen were evaluated for motif enrichment in cattle, pig, and mouse. Adipose- and hypothalamus-specific peaks were excluded from this analysis, due to the low number of tissue-specific peaks detected, and lack of hypothalamus data in the mouse. Several TF families demonstrated consistent motif enrichment in particular tissues, such as forkhead box family members, which were enriched in liver- and lung-specific open chromatin in all three species (Figure 3d). Homeobox motifs also demonstrated consistent enrichment patterns in tissue-specific open chromatin across species; HNF motifs were enriched in liver, NKX3.1 motifs in lung, and MEIS1 and SIX1 motifs in muscle. Brain-specific open chromatin was consistently enriched for motifs of brain-specific TFs (ATOH1, NEUROD1, and OLIG2). Nevertheless, several discrepancies between species were noted in motif enrichment of tissue-specific open chromatin. For instance, bHLH factors MYF5, MYOD, and MYOG motifs were consistently enriched in muscle-specific open chromatin, but only demonstrated enrichment in cortex-specific open chromatin in the mouse. Across species, spleen-specific open chromatin demonstrated little motif enrichment, with the notable exception of ETS, IRF, and Runt TF family motifs, which were almost exclusively enriched in the pig. Overall, liver-, lung-, and muscle-specific regulatory circuitry appeared to be the most highly conserved across cattle, pig and mouse, whereas brain-specific regulation was more varied between species.

### Conservation of chromatin accessibility across mammals

Although extensive epigenetic divergence is expected between species, sequence similarity among cattle, pig, and mouse genomes well above the coding sequence fraction suggests that a significant portion of epigenetic control of transcription is likely under evolutionary constraint. Having observed similarities in motif enrichment in tissue-specific open chromatin between species, it was suspected that the sequence and accessibility of regulatory elements would also be constrained.

Reasonably, portions of regulatory elements, such as the motifs that facilitate TF-DNA contacts, would be under selective pressure, especially those that connect tissue-specific transcription factors to conserved tissue-specific expression programs. To identify homologous regions that correspond to regulatory elements, the coordinates of consensus ATAC-seq peaks from each species were projected to the other two species using Ensembl Compara, which is largely based on whole-genome pairwise and multiple sequence alignments. Unsurprisingly, considering the smaller size of the mouse genome and the larger relative evolutionary distance between mice and ungulates, more consensus peaks could be mapped between pig and cattle, as opposed to between pig and mouse or cattle and mouse. Overall, about half of peaks were conserved at the sequence level between cattle and pig, whereas only about a third of peaks were conserved at the sequence level between ungulates and mice (Figure 4a). Moreover, about 40% of accessible regions that could be mapped between pig to cattle were accessible in at least one tissue in both species, whereas only about 30% of accessible regions mapped between mouse and pig, or between mouse and cattle, demonstrated conserved accessibility in at least one tissue (Figure 4a; Figure S6). Overall, conservation of open chromatin at specific loci was in line with evolutionary distance. Comparing cattle and pig, which are separated by about 62 million years, about 20% of consensus peaks were conserved in terms of sequence and accessibility, whereas mouse, separated from ungulates by about 96 million years, only shared about 10% of consensus peaks, in terms of sequences and accessibility, with either species (Figure S7a,b). Additionally, promoter accessibility at homologous regions was considerably more conserved than enhancer accessibility at homologous regions, with almost half of promoter open chromatin in the pig detected in cattle, while only a fifth of all open chromatin in pig was conserved in cattle (Figure S7a,b).

**Figure 4.**
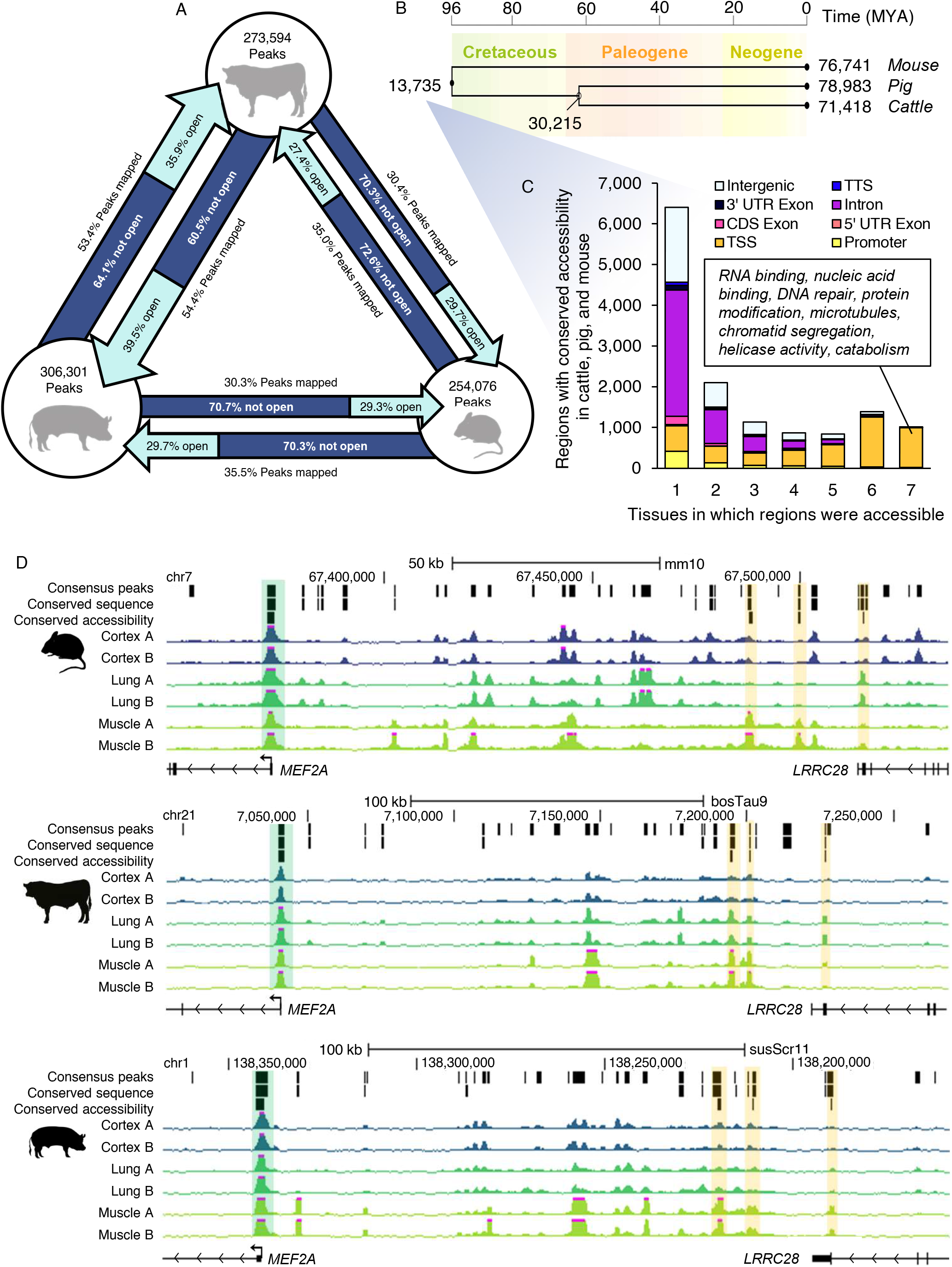
Conservation of regulatory element accessibility in cattle, pig, and mouse. A) Summary of pairwise open chromatin conservation. Circles reflect number of consensus ATAC-seq peaks in each species. Arrow width reflects proportion of consensus peaks that could be projected to the other species. Lighter section of arrows reflect proportion of regions that could be mapped which demonstrated conserved accessibility in at least one tissue in both species. B) Of regions that could be projected to all three species, number of regions with conserved accessibility in all three species and in ungulates laid over a phylogenetic tree reflective of evolutionary distance in millions of years (MYA). C) Genomic distribution of regions with conserved accessibility in all three species, relative to the mouse gene annotation. Brief summary of enriched gene ontology (GO) terms for genes marked by conserved open chromatin at their TSS in every tissue. D) Consensus peaks, consensus peaks with conserved sequence (that could be mapped to all three species), and consensus peaks with conserved sequence and conserved accessibility in all three species at the MEF2A locus. Tracks show normalized ATAC-seq signal. Conserved promoter open chromatin highlighted in green. Conserved distal (putative enhancer) open chromatin highlighted in yellow.

Among the consensus peaks identified in cattle, pig and mouse, 145,801 regions could be mapped to all three species. Of these, 13,735 were consistently accessible in the same tissue (at least one) in all three species, and 30,215 were accessible in at least one tissue in both cattle and pig (Figure 4b). Regions with conserved accessibility in all three species tended to be ubiquitously present in all sampled tissues. Whereas only 2-4% of consensus peaks were accessible in all tissues when considering a single species, 7% of regions with conserved accessibility in all species were accessible in all tissues (Figure S7c). Furthermore, regions with conserved accessibility in all three species were heavily enriched around TSS (32% of regions), especially those which were accessible in all tissues (97% of regions), which marked TSS of housekeeping genes (Figure 4c; Table S2). In contrast, regions that demonstrated conserved accessibility in cattle and pig, but not in mouse, were very rarely accessible in all tissues (only 23 out of 14,543 regions, or 0.2%) (Figure S7d), and only occurred at TSS 4% of the time (Figure S7e).

Intriguingly, most regions with conserved accessibility in cattle, pig and mouse were only open in one or two tissues (62%) and were predominantly intronic and intergenic (Figure S7e). For instance, several regions upstream of the *MEF2A* locus demonstrated muscle-specific accessibility in all three species (Figure 4d). These regions could represent conserved enhancers, suggesting that even distal regulatory elements are subject to some level of evolutionary constraint. In all, 3,105 intergenic loci (6%) demonstrated conserved open chromatin signatures in all three species, and further examination of the genes that were closest to these sites (within 100 kb) revealed functional enrichment for developmental processes, such as regionalization and organogenesis (Table S3).

Notably, this small subset of “conserved” enhancers may underrepresent conserved activity at distal loci. For instance, accessible regions around the FOXG1 locus appear to be syntenically conserved across all three species; however, only one region – corresponding to the TSS of a human long non-coding RNA – could be mapped to all three species (Figure S8). The remaining loci could not be mapped between two species, suggesting a pervasive lack of overall sequence identity, despite apparent functional conservation.

## Discussion

Despite the intimate connection between chromatin structure and regulation of transcription, an atlas of chromatin accessibility in livestock tissues had not yet been reported. To address this gap in knowledge, ATAC-seq was used to identify regions of open chromatin in a prioritized set of pig and cattle tissues, yielding a first glimpse at the landscape of active regulatory elements in these genomes. In all, 6 to 7% of the cattle and pig genomes demonstrated accessibility in at least one tissue, which was consistent with a comparable dataset in the mouse (26). Notably, about half of these accessible sites were intergenic, accounting for about 3% of each genome. The identification of these regulatory elements is a crucial first step towards a comprehensive annotation of the non-coding genome, which has been severely lacking in livestock species (37).

Although efforts are currently underway to further characterize these regulatory elements as enhancers, silencers, insulators, promoters, etc. (37,38), motif enrichment analysis highlighted some of their potential regulatory roles. By far, the most enriched sequences in cattle and pig open chromatin were CTCF motifs, suggesting pervasive involvement of accessible regions in higher order chromatin organization. In particular, convergently oriented CTCF sites are known to demarcate topologically associated domain (TAD) boundaries (39), which are largely invariable across cell types (40–42) and even across species (40,41,43,44). Indeed, regions that were globally accessible in all pig and cattle tissues were particularly enriched for CTCF recognition motifs, suggesting that these regions may delineate TAD boundaries, although direct profiling of chromatin interactions will be necessary to provide more definitive annotations of 3D chromatin structure.

Interestingly, out of all accessible CTCF motifs, almost a third were only accessible in a single tissue, which is consistent with CTCF binding being highly variable in different cell types (45,46), even though TAD structure is largely consistent (40–42). Only a fraction of CTCF binding sites (15%) actually localize to TAD boundaries (47), and most CTCF binding sites are interspersed with enhancers, stabilizing enhancer-promoter interactions (48), and forming cell type-specific chromatin loops linked to differential gene expression (48–50). Therefore, differentially accessible CTCF motifs may reflect tissue-specific chromatin looping. Taken together, the presence of both tissue-specific and globally accessible CTCF motifs suggests a multi-tiered 3D structure that participates in both fundamental and tissue-specific regulation.

Tissue-specific open chromatin was widespread and conspicuously lacking near TSS. Motif enrichment analysis revealed that tissue-specific open chromatin demonstrated conserved enrichment for tissue-specific TF binding motifs, which was expected, as expression programs in vertebrate tissues are thought to be controlled by a highly conserved set of tissue-specific TFs (51). However, not all TF families demonstrated consistent motif enrichment in tissue-specific open chromatin. The motifs of the RUNX family, highly conserved TFs involved in cell fate determination (52), were only enriched in mouse lung-, pig cerebellum-, and spleen-specific open chromatin. Whether this points to a divergence in tissue-specific regulation remains unclear, as motif enrichment analyses rely heavily on known binding motifs in human and mouse, failing to account for any species-specific differences in TF recognition sites.

Certainly, divergent chromatin structure has implications for differential transcriptional regulation, a phenomenon that has been long recognized as a significant contributor to phenotypic diversity (34,35,53–55). As expected, the proportion of open chromatin that was conserved between species was consistent with evolutionary distance – higher concordance was observed between cattle and pig, than between mouse and either cattle or pig. By classifying loci as either proximal or distal based on their closeness to annotated genes, it was also apparent that accessibility at proximal elements, such as promoters, was significantly more conserved than accessibility at distal elements. This discrepancy in functional conservation was not altogether surprising; whereas promoters are fundamental for gene expression in any context, modulation of enhancer activity can subtly alter phenotypes without compromising viability (56,57). In fact, enhancers are known to evolve rapidly, and several studies have demonstrated how changes to enhancer sequences can lead to differing phenotypes between species (36,58–60). Indeed, only 17% of intergenic open chromatin in cattle was also accessible in pig, and a meager 6% was accessible in mice, indicating that enhancers are largely species-specific, as has been previously demonstrated (3,35). Nevertheless, more than 3,000 intergenic loci, relative to gene annotations in the mouse, had a conserved open chromatin signature in at least one tissue in all three species. Considering some highly conserved enhancers have been implicated in core biological processes, such as embryonic development (33), these intergenic loci are suspected to be involved in fundamental biological processes in adult tissues, which would account for their abnormal sequence constraint and functional conservation.

Intriguingly, several loci appeared to share open chromatin signatures based on synteny, despite lack of sequence conservation. Several studies have demonstrated that enhancer function can be conserved even when overall sequence is not (61–63). Instead, selective pressure may only operate on the functional components of regulatory elements: TF binding sites, which are typically short and degenerate sequences (34–36). Although inferring sequence conservation is possible with sequences as short as 36 bp (34), detecting homologous regulatory regions based only on conserved TF binding sites (6-12 bp (64)) is problematic. This begs a pragmatic question in the field of comparative epigenomics: if orthologous regions cannot be determined based on sequence, how then can we determine whether function is conserved? If TF binding sites are all that is required for enhancer function, then most enhancers would not be conserved in the canonical sense of sequence constraint, but instead through TF binding and relative positioning.

## Conclusions

To our knowledge, these data constitute the first atlas of chromatin accessibility in a common set of livestock tissues, and consequently a first look at the distribution across multiple tissues of active regulatory elements in the pig and cattle genomes. Moreover, this initial annotation of the non-coding genome will help to inform the identification of causal variants for disease and production traits. From the standpoint of comparative epigenomics, these data contribute to the ever-growing wealth of epigenomic information; the comprehensive analysis of which will undoubtedly help bridge the gap between genome and phenome, providing crucial insight into transcriptional regulation and its connection to evolution.

## Methods

### Tissue collection and cryopreservation

All necessary permissions were obtained for collection of tissues relevant to this study, following the Protocol for Animal Care and Use #18464, as per the University of California Davis Animal Care and Use Committee (IACUC). As described previously (65), two intact male Line 1 Herefords, provided by Fort Keogh Livestock and Range Research lab, were euthanized by captive bolt under USDA inspection at the University of California, Davis. Both cattle were 14 months of age and shared the same sire. Two castrated male Yorkshire pigs were humanely euthanized by animal electrocution followed by exsanguination, which is the standard method of euthanasia at pig slaughterhouses, under USDA inspection at Michigan State University Meat Lab. Pigs were littermates aged six months old, and sourced from the Michigan State University Swine Teaching and Research Center. From each animal, subcutaneous adipose, frontal cortex, cerebellum, hypothalamus, liver, lung (left lobe), longissimus dorsi muscle, and spleen were collected and promptly processed for cryopreservation. For each sample, roughly one gram of fresh tissue was minced and transferred to 10 mL of ice-cold sucrose buffer (250 mM D-Sucrose, 10 mM Tris-HCl (pH 7.5), 1 mM MgCl_2_; 1 protease inhibitor tablet per 50 mL solution just prior to use). Minced tissue was twice homogenized using the gentleMACS dissociator “E.01c Tube” program. Homogenate was filtered with the 100 μm Steriflip vacuum filter system, volume was brought up to 9.9 mL with sucrose buffer, and 1.1 mL DMSO was added to achieve a 10% final concentration. Preparations were aliquoted into cryovials and frozen at −80°C overnight in Nalgene Cryo 1°C freezing containers, then stored at −80°C long-term.

### ATAC-seq library construction and sequencing

A modified ATAC-seq protocol compatible with cryopreserved tissue samples was employed (25). Cryopreserved tissue samples were thawed on ice, then centrifuged for 5 min at 500 rcf and 4°C in a centrifuge with a swinging bucket rotor. Pellets were resuspended in 1 mL ice-cold PBS, and centrifuged again for 5 min at 500 rcf and 4°C. Pellets were then resuspended in 1 mL ice-cold freshly-made ATAC-seq cell lysis buffer (10 mM Tris-HCl pH=7.4, 10 mM NaCl, 3 mM MgCl_2_, 0.1% (v/v) IGEPAL CA-630), and centrifuged for 10 min at 500 rcf and 4°C. Pellets were then resuspended again in ice-cold PBS for cell counting on a hemocytometer. Between 50,000 and 1,000,000 cells were aliquoted for library preparation, depending on tissue, success of previous library preparation attempts, and cell abundance in a given preparation (Table S1). Aliquoted cells were centrifuged once more for 5 min at 500 rcf and 4°C, and pellets were resuspended in 50 μL transposition mix (22.5 μL nuclease-free H_2_O, 25 μL TD buffer and 2.5 μL TDE1 enzyme from Nextera DNA Library Prep Kit (Illumina, cat. no. FC-121-1030)). Nuclear pellets were incubated with transposition mix for 60 min at 37°C, shaking at 300 rpm. Transposed DNA was purified with the MinElute PCR Purification Kit (Qiagen, cat. no. 28004) and eluted in 10 μL Buffer EB. Eluted DNA was added to 40 μL PCR master mix (25.4 μL SsoFast™ EvaGreen® Supermix, 13 μL nuclease-free H_2_O, 0.8 μL 25 μM Primer 1, 0.8 25 μM Primer 2 (see Table S4 for sequences)) to 10 μL eluted DNA and PCR cycled (1 × [5 min at 72°C, 30 sec at 98°C], 10-13x [10 sec at 98°C, 30 sec at 63°C, 1 min at 72°C]). Libraries were then purified again with the MinElute PCR Purification Kit, and eluted in 10 μL Buffer EB. Libraries were quantified by Qubit (Thermo Fisher Scientific, Inc., Waltham, MA), and checked for nucleosomal laddering using a Bioanalyzer High Sensitivity DNA Chip (Agilent Technologies, Santa Clara, CA) (Figures S9 and S10). Details for individual library preparations, including cell input, PCR cycles, and concentrations can be found in Table S1. Finally, libraries were size selected for subnucleosomal length fragments (150-250 bp) on the PippinHT system using a 3% agarose cassette (Sage Science, Beverly, MA). Size selection and DNA concentration were evaluated with a Bioanalyzer High Sensitivity DNA Chip, pooled, and submitted for sequencing on the NextSeq 500 platform to generate 40 bp paired end reads.

### ATAC-seq data preprocessing and quality evaluation

Low quality bases and residual adapter sequences were trimmed from raw sequencing data using Trim Galore! (v0.4.0), a wrapper around Cutadapt (v1.12)(66), with the options “-a CTGTCTCTTATA” and “-length 10” to retain trimmed reads at least 10bp in length. Trimmed reads were then aligned to either the susScrofa11 (pig), ARS-UCD1.2 (cattle), or GRCm38 (mouse) genome assemblies using BWA mem with default settings (v0.7.17)(67). Duplicate alignments were removed with Picard-Tools (v2 .9.1), and mitochondrial and low quality (q < 15) alignments were removed using SAMtools (v1.9)(68). Finally, broad peaks were called using MACS2 (v2.1.1)(69) with options “-q 0.05 -B –broad –nomodel –shift −100 –extsize 200.”

For all reported statistics, standard error is also reported. Quality metrics were calculated as follows. The non-redundant read fraction (NRF) was calculated by dividing the number of nonduplicate uniquely mapping reads out of all mapped reads. The Fraction of Reads in Peaks (FRiP) score for each sample was calculated using the plotEnrichment function from the deepTools suite (v.3.2.0), given the broad peaks identified by MACS2. Finally, the synthetic Jensen-Shannon distance (sJSD) was calculated using the deepTools plotFingerprint function.

For visualization, genome-wide ATAC-seq signal was normalized by RPKM in 50 bp windows using the bamCoverage function from the deepTools suite. Other deepTools functions were used along with the resulting bigwig files to generate (1) pairwise scatter plots of genome-wide signal, including Spearman correlation coefficients (plotCorrelation –log1p – removeOutliers), (2) principal components analyses of signal in consensus peaks (plotPCA – transpose –log2), and signal at loci of interest (plotHeatmap), such as TSS (computeMatrix reference-point –beforeRegionStartLength 2000 –afterRegionStartLength 2000 –skipZeros) and peaks (computeMatrix scale-regions –beforeRegionStartLength 500 –regionBodyLength 500 – afterRegionStartLength 500 –skipZeros). Normalized signal from bigWig files was visualized at specific loci with the UCSC Genome Browser, limiting the y-axis range to 20 and displaying the mean in smoothing windows of 4 bases.

### Identification of consensus and tissue-specific open chromatin

Broad peaks called from biological replicates in each tissue were compared using BEDtools intersect (70) (v2.26.0) to identify consensus peaks. In the case of cattle cerebellum, for which only one biological replicate was available, consensus peaks were identified by calling broad peaks more stringently (-q 0.01 -B –broad –nomodel –shift −100 –extsize 200). Tissue-specific open chromatin was defined as consensus peaks that were not detected as accessible in either biological replicate of any other tissue.

### Categorization of peaks by location relative to gene annotations

Peaks were categorized by position relative to features in the Ensembl annotation (v96), using BEDtools intersect. Because many peaks overlap multiple features, peaks were first classified as TSS (within 50bp), then as promoters (within 2kb upstream of TSS), as transcription termination sites (TTS; within 50 bp), as overlapping a 5’ untranslated region (UTR), as overlapping a 3’ UTR, as exonic, as intronic, and finally, if peaks did not overlap any of these features, they were considered to be intergenic. The locations of peak sets were randomized using BEDtools shuffle, and these were also categorized by location relative to annotated genes, such that the proportion of peaks localizing to any given genomic feature could be compared to the expected proportion given random genomic intervals.

### Motif enrichment analysis

Regions of interest were evaluated for motif enrichment using the HOMER findMotifs.pl function (v4.8) (71), and the top ten enriched known motifs, based on p-values, were reported.

### Conservation of open chromatin

All interspecies comparisons were based on the 46-mammalian Enredo-Pecan-Ortheus (EPO) multiple sequence alignment (MSA) available through Ensembl Compara (v99) (32). Regions of consensus open chromatin in each species were projected onto the other two species using the Ensembl Compara Application Programming Interface (API). For simplicity, regions that mapped to multiple loci in another species were discarded prior to evaluating whether accessibility was conserved. Chromatin accessibility was considered to be conserved at homologous regions if they overlapped (by at least 1 bp) consensus open chromatin in the same tissue (at least one tissue) in all species in question.

### Functional annotation enrichment analysis

Ensembl IDs were converted to external gene names using the BiomaRt package, and these were submitted to DAVID (v6.8) (72,73) for functional annotation clustering. Mus musculus was used as background, and functional annotation clustering was conducted on medium stringency for the following terms: GOTERM_BP_5, GOTERM_CC_DIRECT, GOTERM_MF_DIRECT, BIOCARTA, and KEGG_PATHWAY. For each gene set, the top four clusters were reported.

## Supporting information

Supplementary Data 1. Cattle ATAC-seq peaks.

Supplementary Data 2. Pig ATAC-seq peaks.

## Declarations

## Abbreviations

ATAC-seq: Assay for Transposase Accessible Chromatin
CDS: Coding sequence
ChIP-seq: Chromatin immunoprecipitation followed by sequencing
DNase-seq: DNase I hypersensitive sites sequencing
FRiP: Fraction of reads in peaks
GO: Gene ontology
MYA: Million years ago
NRF: Non-redundant read fraction
PCA: Principal components analysis
RPM: Reads per million
sJSD: Synthetic Jensen-Shannon distance
TAD: Topologically associated domain
TF: Transcription factor
TSS: Transcription start site
TTS: Transcription termination site
UTR: Untranslated region

## Ethics approval and consent to participate

All necessary permissions were obtained for collection of tissues relevant to this study, following the Protocol for Animal Care and Use #18464, as per the University of California Davis Animal Care and Use Committee (IACUC).

## Consent for publication

Not applicable.

## Availability of data and materials

ATAC-seq data generated by this study is available from the European Nucleotide Archive under project ID PRJEB14330 (https://www.ebi.ac.uk/ena/data/view/PRJEB14330). Raw ATAC-seq data for male mesenteric fat, cerebrum, cerebellum, liver, lung, skeletal muscle and spleen from mice (26) were downloaded from the CNGB Nucleotide Sequence Read Archive, under Project ID CNP0000198.

## Authors’ contributions

MMH, HZ and PJR designed the study. JM, AVE, IK, CKT, CWE contributed to the experimental design. MMH, PS, YW, and GC collected samples and constructed ATAC-seq libraries. MMH conducted bioinformatics analyses with guidance from CK. MMH, HZ and PJR wrote the manuscript. All authors have read and approved the manuscript.

## Acknowledgements

Not applicable.

---

**Figure S1.**
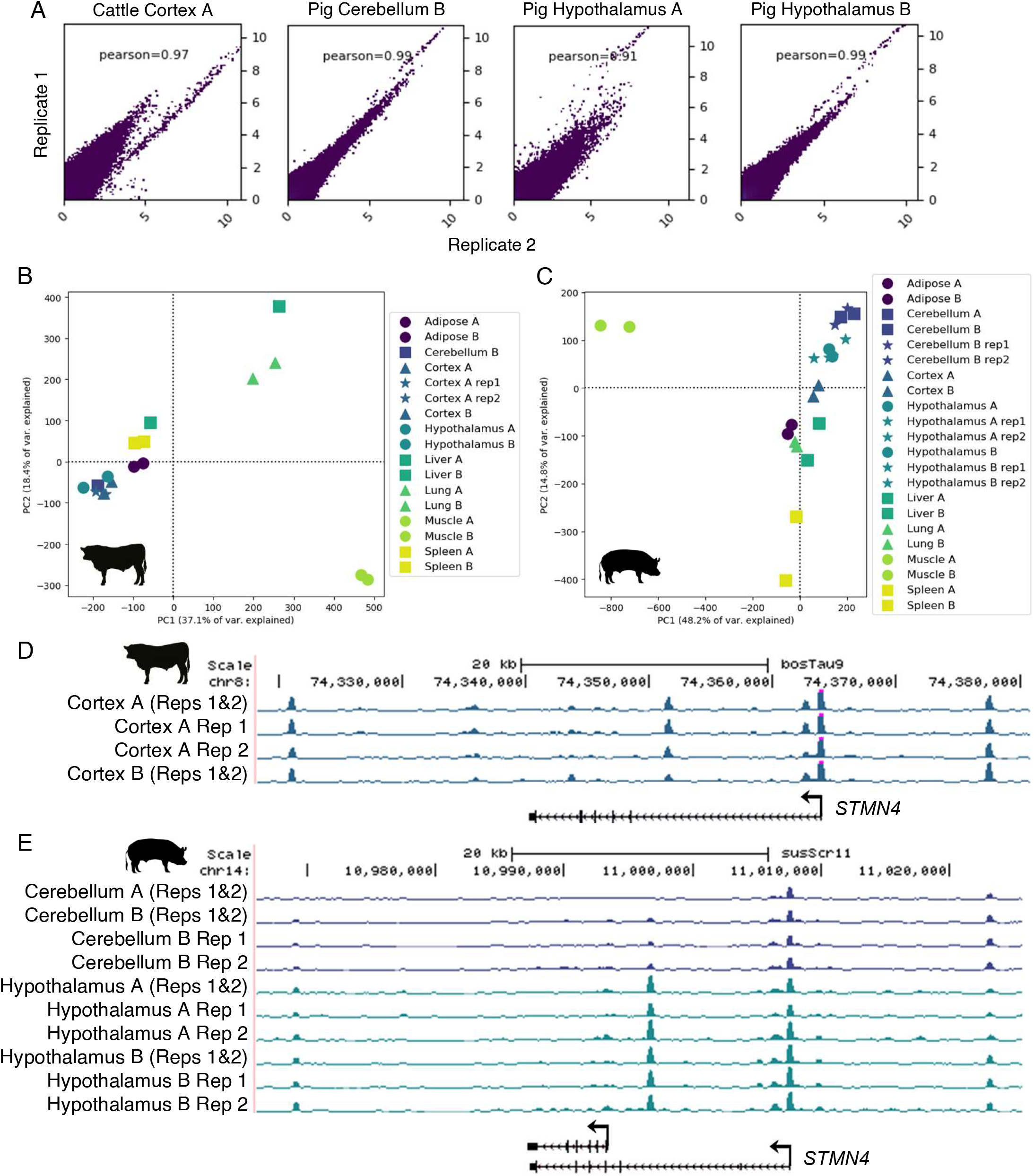
Correlation of ATAC-seq signal in select technical replicates ATAC-seq libraries. A) Pearson correlation of genome-wide signal (RPKM) in 500 bp windows. B) PCA of Cortex A technical replicate libraries alongside all biological replicates. C) PCA of pig technical replicate libraries alongside all biological replicates. D) Signal of cattle cortex technical and biological replicates at the STMN4 locus. E) Signal of pig cerebellum and hypothalamus technical and biological replicates at the STMN4 locus.

**Figure S2.**
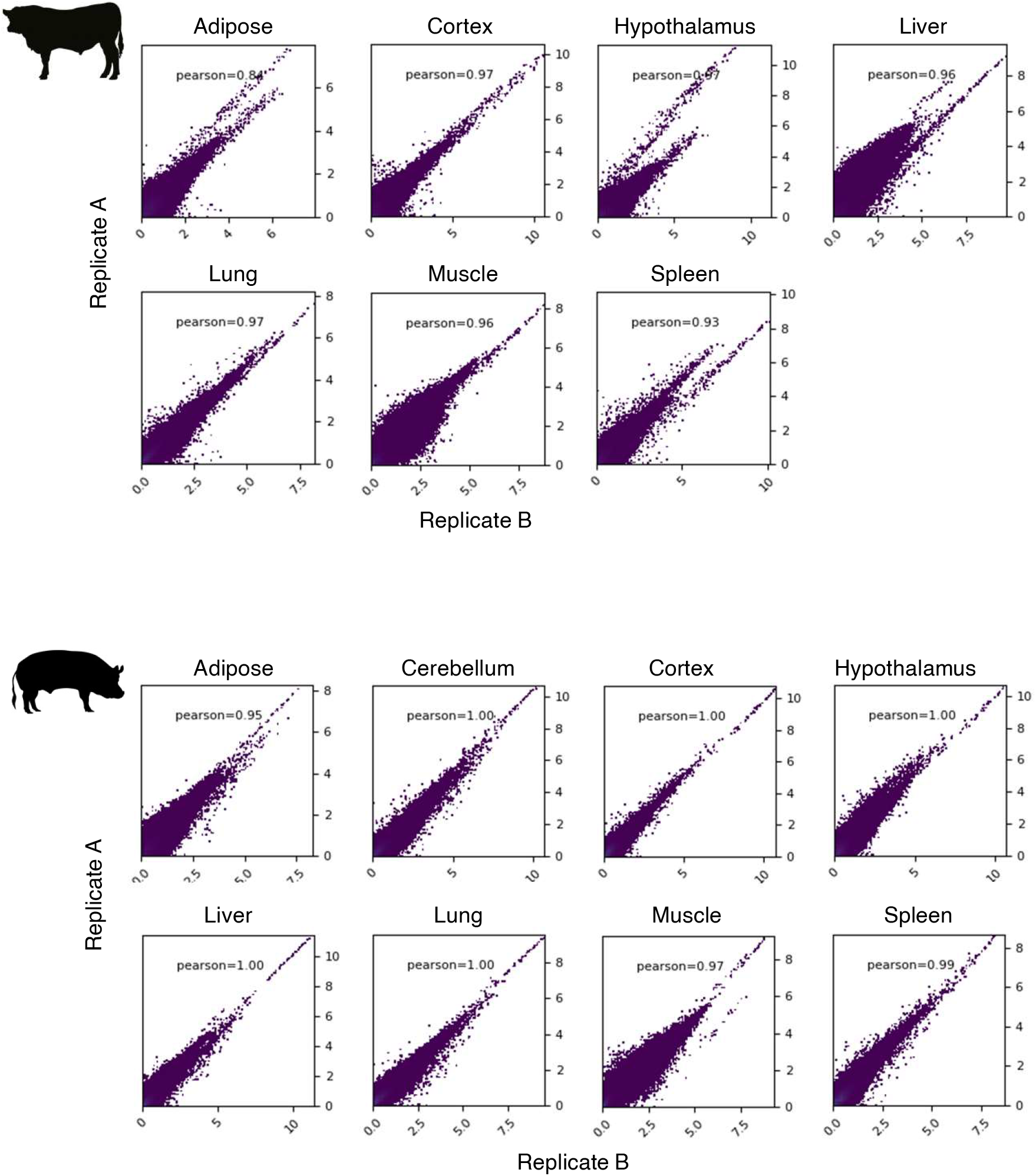
Correlation of ATAC-seq signal in biological replicates. Scatterplots showing Pearson correlation of normalized genome-wide signal in 500 bp windows between biological replicates for cattle and pig tissues.

**Figure S3.**
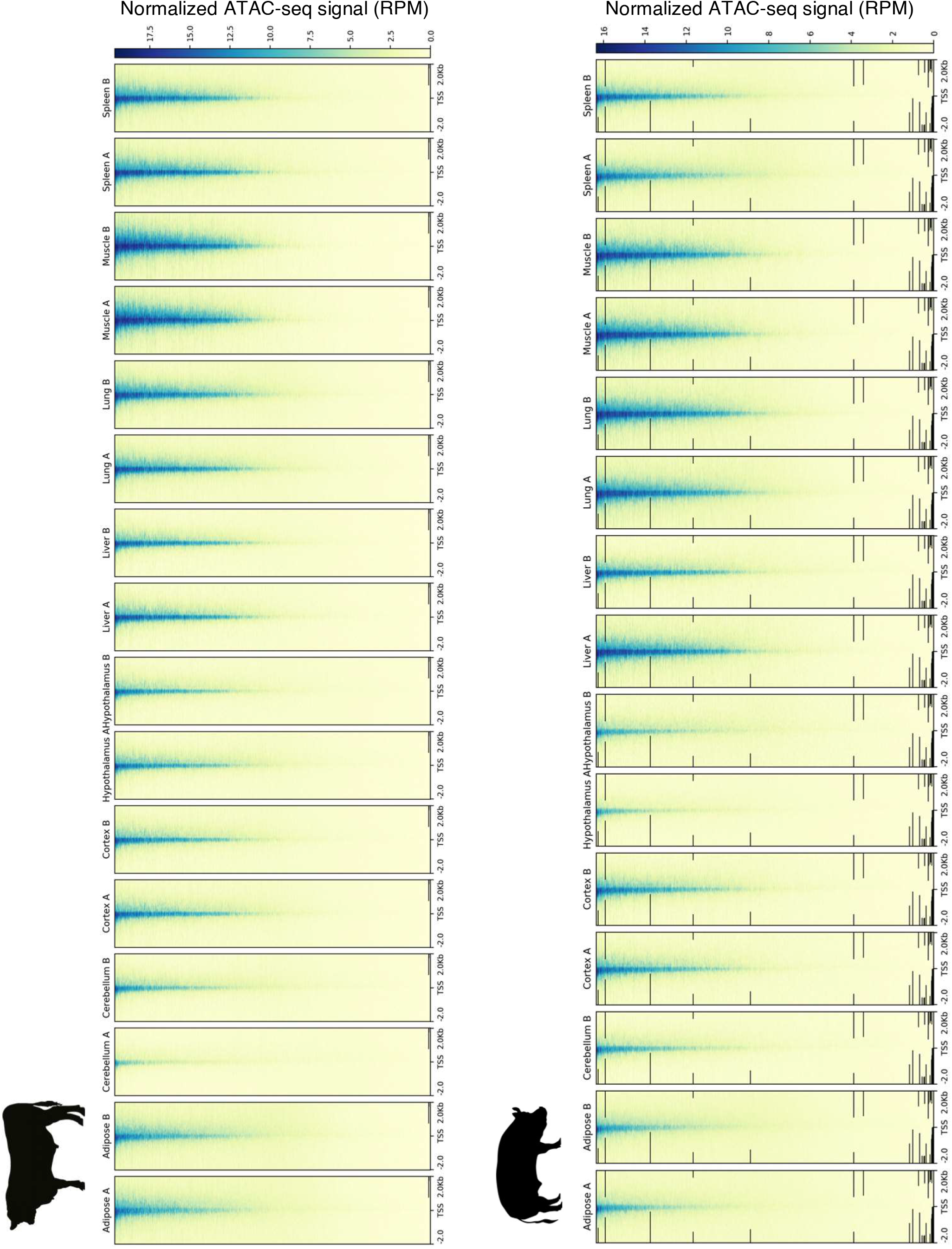
ATAC-seq signal at TSS. Heatmaps depicting normalized ATAC-seq signal in the proximity of TSS, including 2 kb upstream and downstream, with TSS sorted by signal intensity.

**Figure S4.**
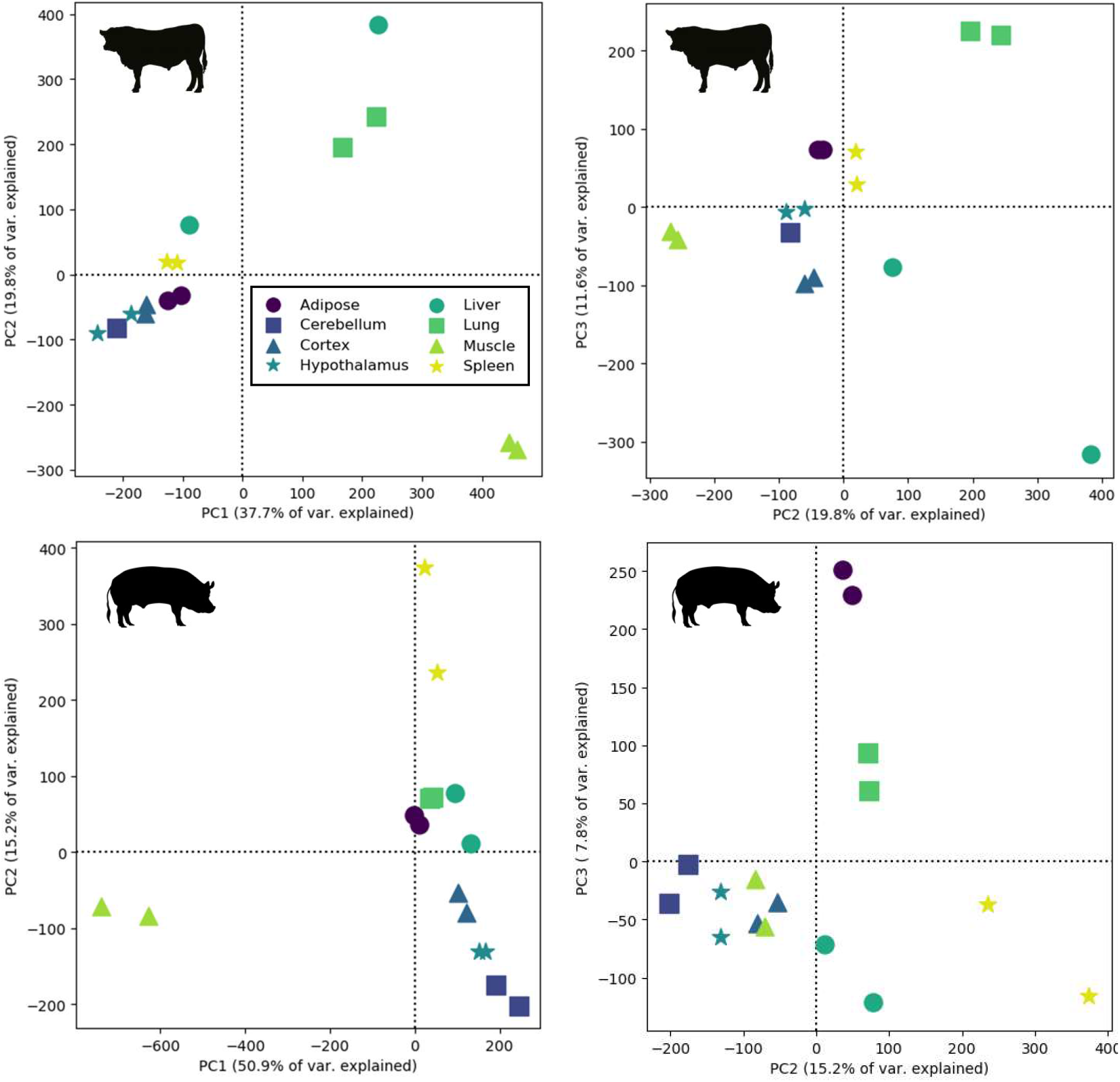
PCA of normalized ATAC-seq signal in consensus open chromatin identified in pig and cattle tissues tissues. Principal components 1, 2 and 3 are included to better visualize clustering of tissues.

**Figure S5.**
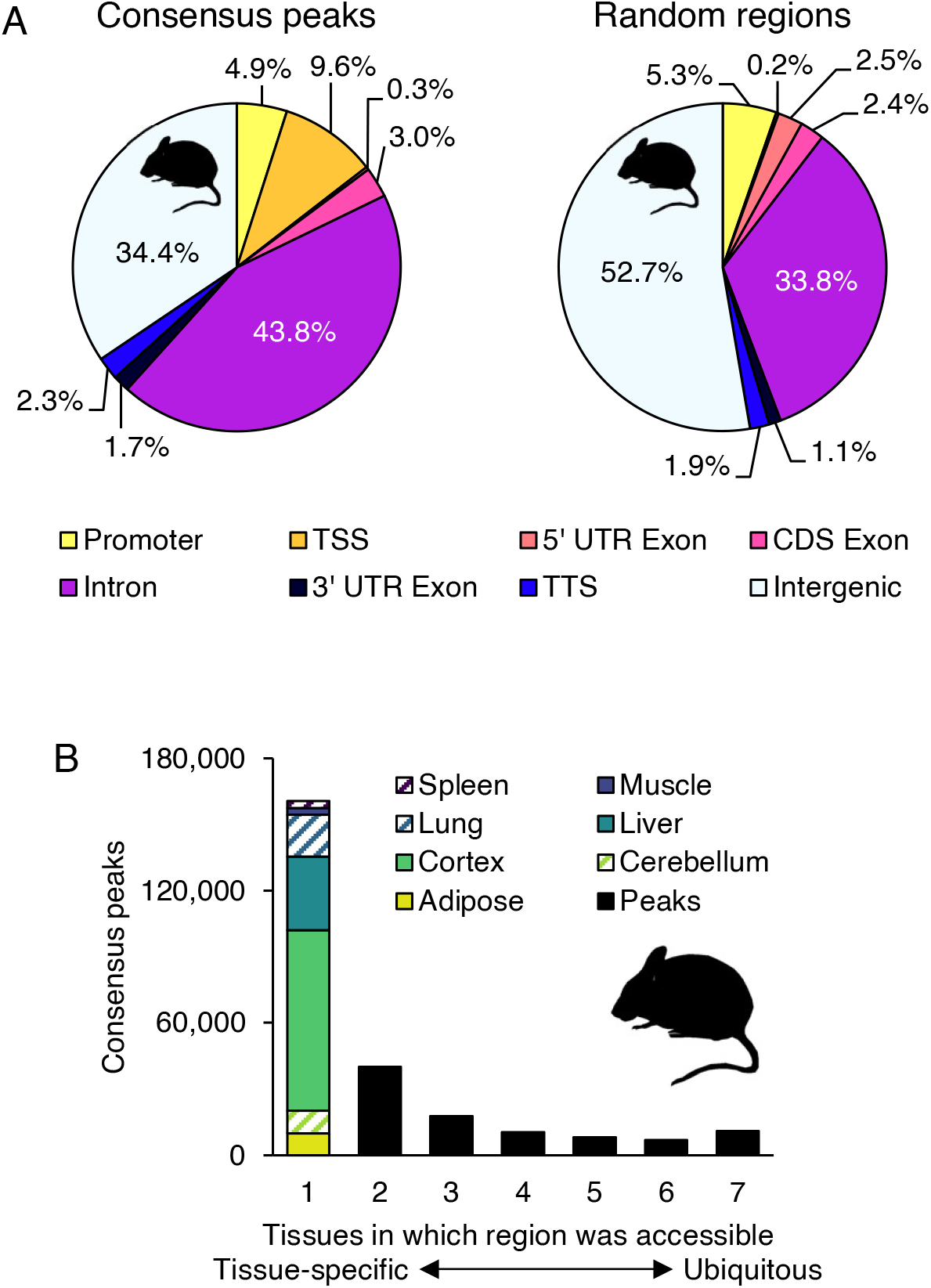
Mouse open chromatin localization and differential accessibility. A) Distribution of mouse consensus open chromatin, as well as randomized genomic intervals, relative to the Ensembl gene annotation (v96). B) Distribution of consensus peak activity, ranging from tissue-specific (accessible in only one tissue) to ubiquitous (accessible in all sampled tissues). Consensus peaks that were accessible in a single tissue were further broken down by tissue.

**Figure S6.**
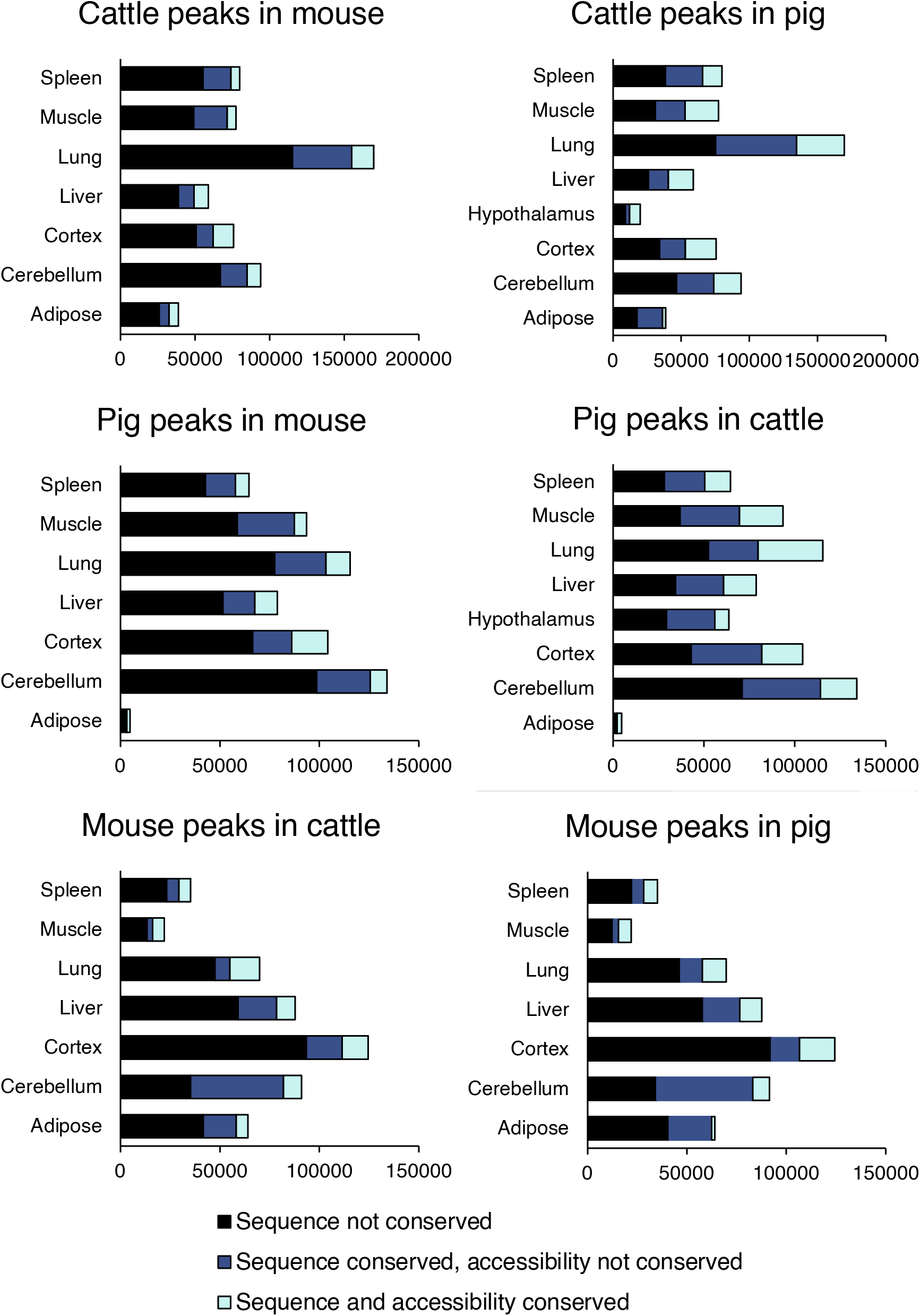
Conservation of open chromatin in individual tissues. Titles above bar plots indicate the species that consensus peaks were identified in, followed by the species to which the consensus peak coordinates were projected to check for sequence and accessibility conservation in the corresponding tissue.

**Figure S7.**
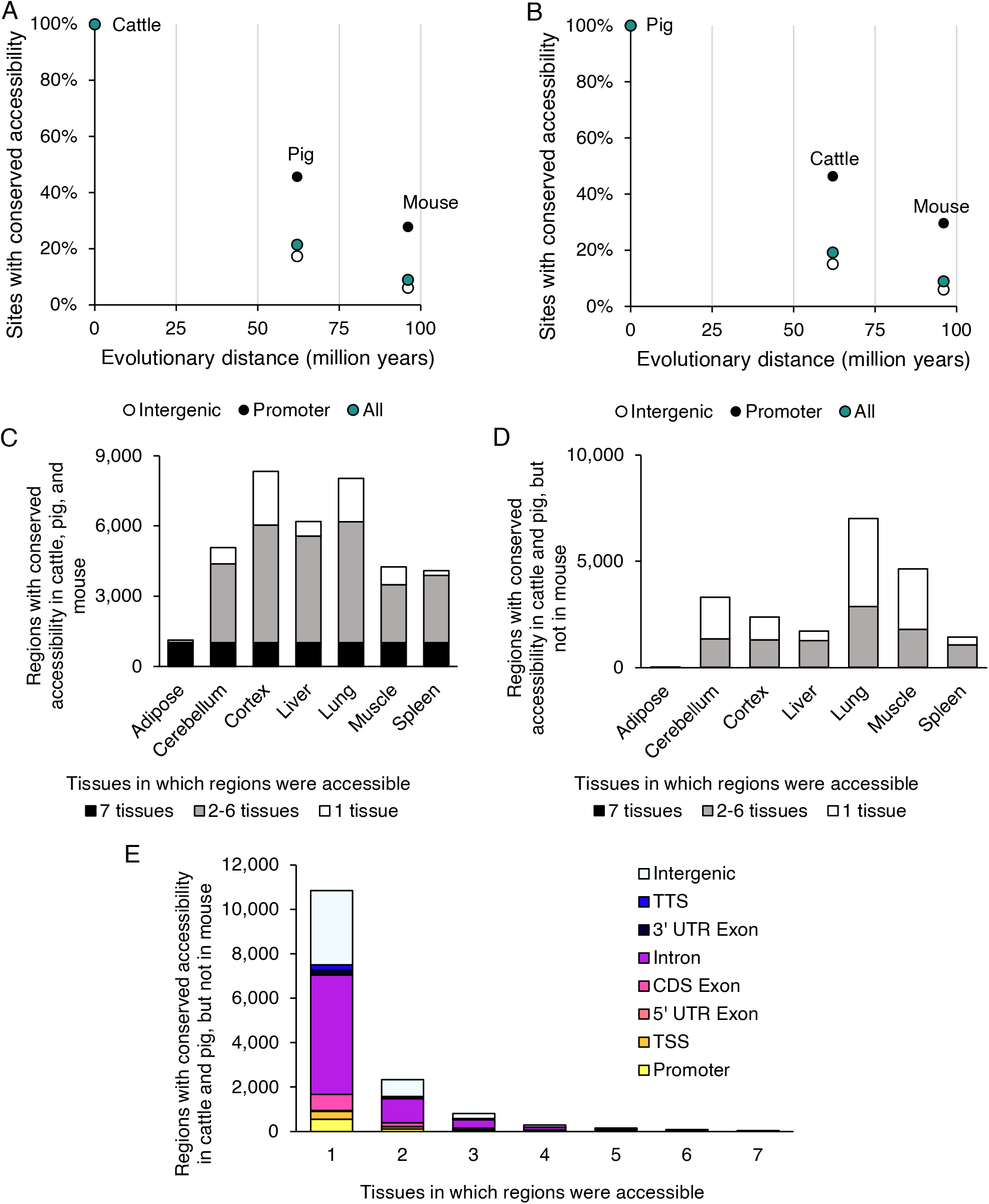
Characterization of conserved open chromatin. Proportion of all consensus peaks, promoter consensus peaks (within 2kb upstream and 50 bp downstream of TSS), and intergenic consensus peaks that were identified in (A) cattle or (B) pig that demonstrated both conserved sequence and accessibility in the other two species. Number of tissues that consensus peaks which were conserved in (C) all three species or (D) only in cattle and pig were accessible in. E) Distribution of consensus peaks with conserved accessibility in cattle, pig, and mouse, relative to the mouse gene annotation.

**Figure S8.**
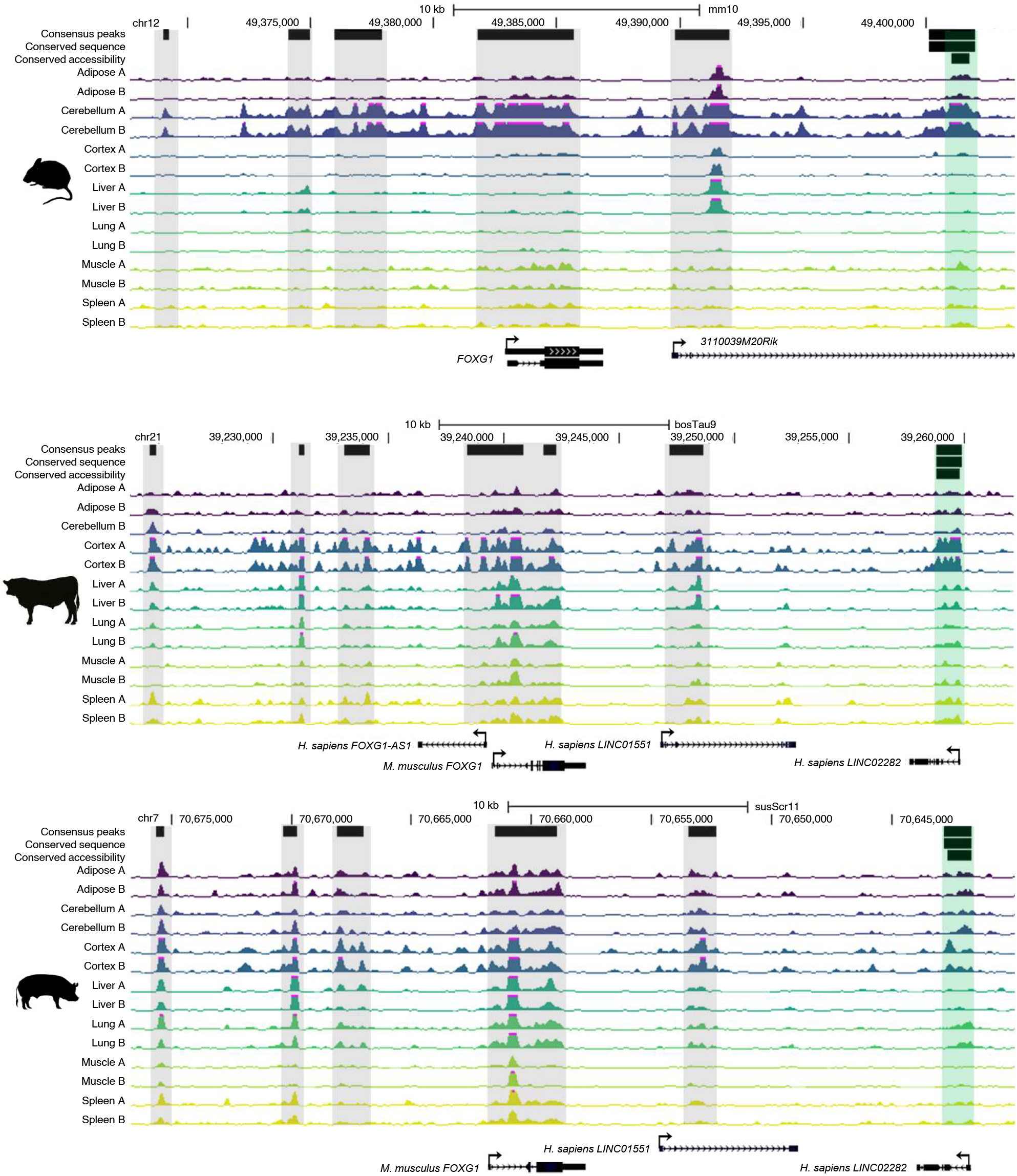
Positional conservation of chromatin accessibility at the *FOXG1* locus. For each species, consensus peaks, consensus peaks with conserved sequence (that could be mapped to all three species), and consensus peaks with conserved accessibility are shown. Tracks show normalized ATAC-seq signal for each sample. Conserved promoter open chromatin is highlighted in green. Consensus peaks that appear to be syntenically conserved, relative to *FOXG1*, but which could not be mapped between species, are highlighted in grey.

**Figure S9.**
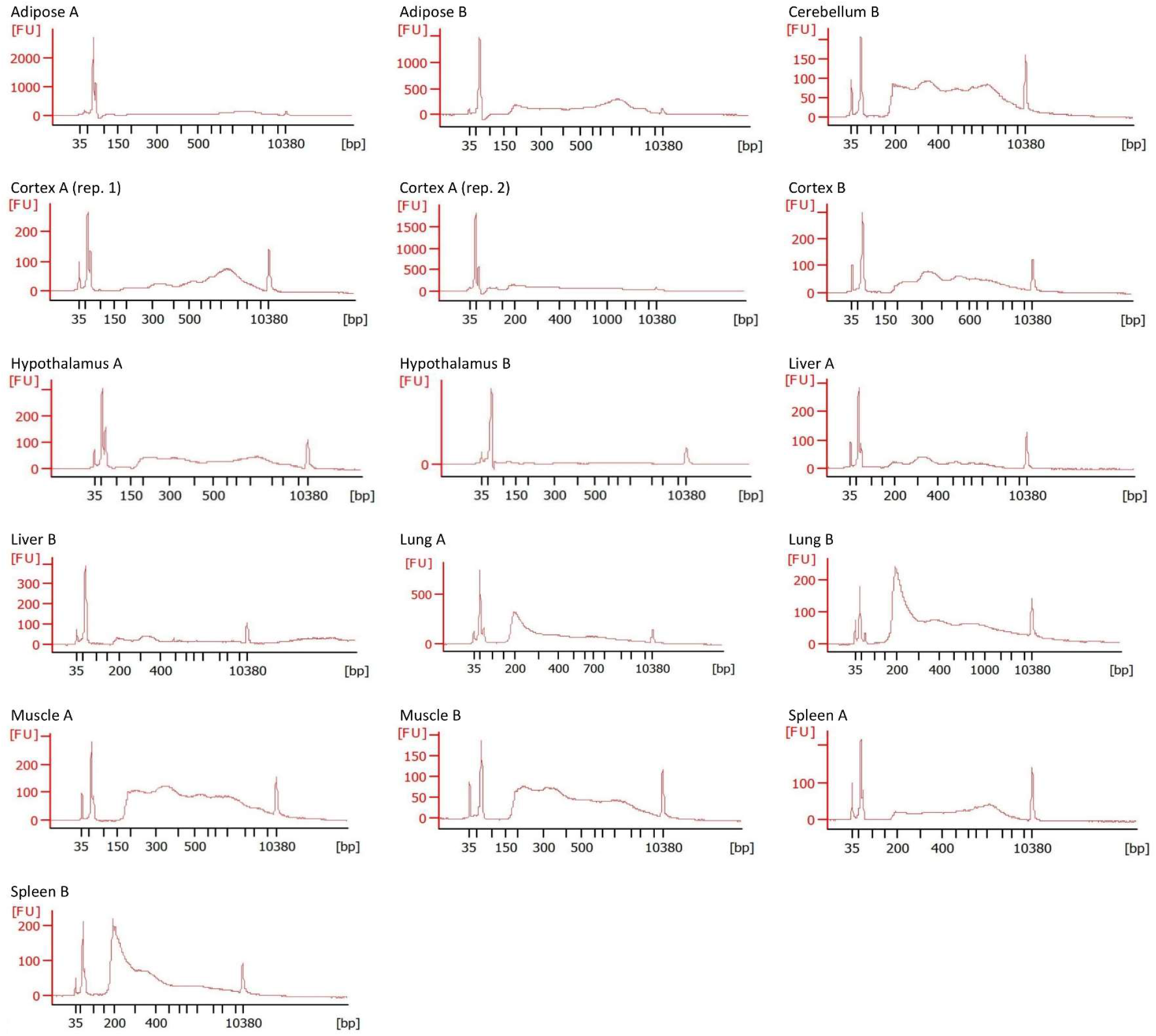
Bioanalyzer traces of cattle ATAC-seq libraries prior to size selection. Bioanalyzer traces were used to check for nucleosomal laddering. Size selection removed excess primer and fragments > 250 bp.

**Figure S10.**
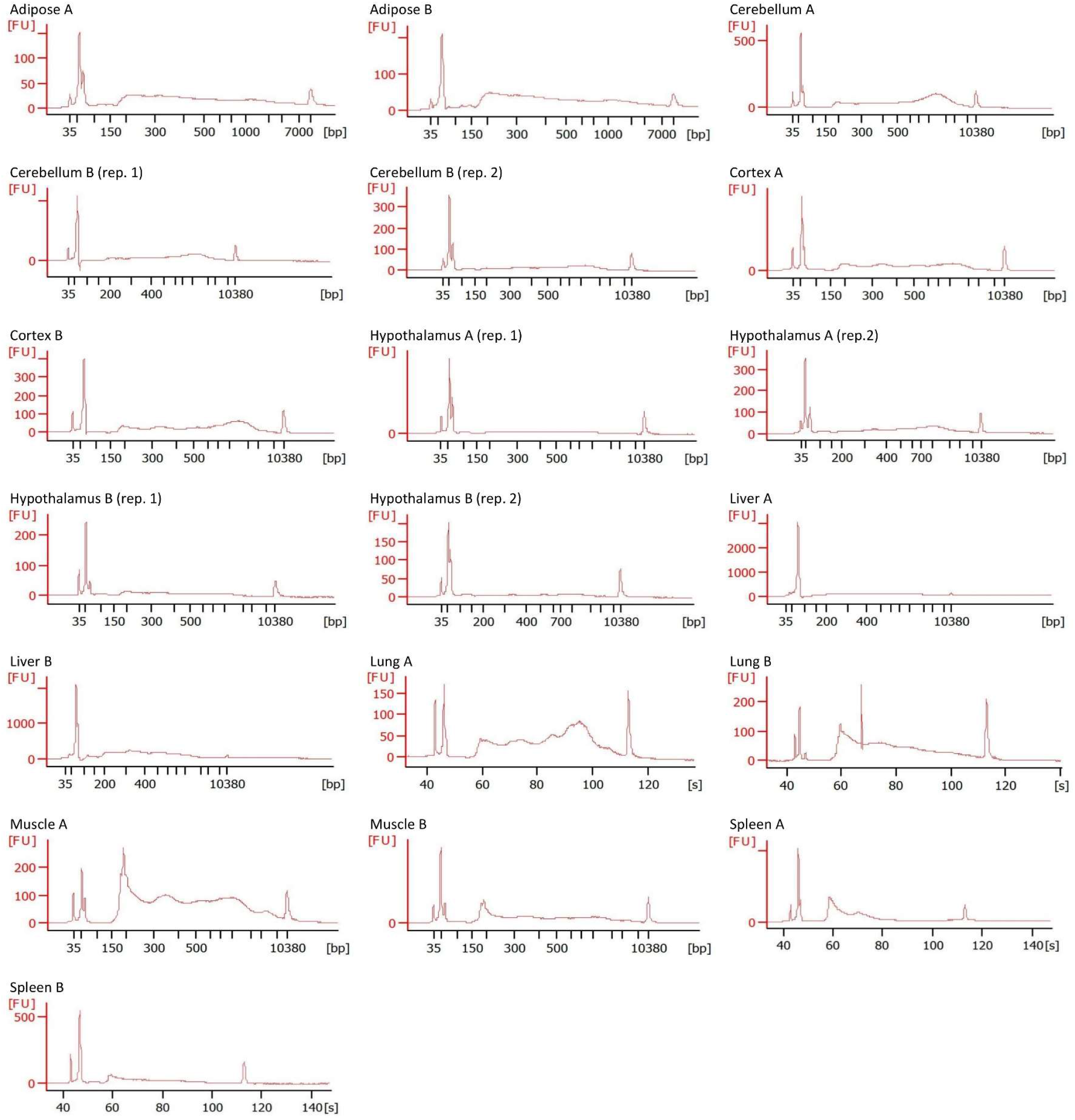
Bioanalyzer traces of pig ATAC-seq libraries prior to size selection. Bioanalyzer traces were used to check for nucleosomal laddering. Size selection removed excess primer and fragments > 250 bp.

**Table S1.** ATAC-seq library construction details. For each library constructed, rounds of PCR amplification, number of cells used as input, and concentration in the 150-250 bp range prior to size selection are indicated.

| Species | Tissue | Biological replicate | Technical replicate | PCR cycles | Cell input | 150-250 bp range |  |
| --- | --- | --- | --- | --- | --- | --- | --- |
|  |  |  |  |  |  | Concentration (ng/ul) | Total DNA (ng) |
| Cattle | Adipose | A | 1 | 10 | 50,000 | 0.51 | 5.08 |
| Cattle | Adipose | B | 1 | 10 | 200,000 | 1.57 | 15.69 |
| Cattle | Cerebellum | B | 1 | 12 | 500,000 | 3.27 | 32.71 |
| Cattle | Cerebellum | B | 1 | 12 | 500,000 | 3.27 | 32.71 |
| Cattle | Cortex | A | 1 | 12 | 200,000 | 0.55 | 5.48 |
| Cattle | Cortex | A | 2 | 13 | 1,000,000 | 2.00 | 19.99 |
| Cattle | Cortex | B | 1 | 12 | 200,000 | 1.57 | 15.71 |
| Cattle | Hypothalamus | A | 1 | 11 | 500,000 | 1.95 | 19.52 |
| Cattle | Hypothalamus | B | 1 | 11 | 500,000 | 0.22 | 2.17 |
| Cattle | Liver | A | 1 | 12 | 50,000 | 1.46 | 14.62 |
| Cattle | Liver | B | 1 | 12 | 500,000 | 1.77 | 17.68 |
| Cattle | Lung | A | 1 | 12 | 50,000 | 12.98 | 129.78 |
| Cattle | Lung | B | 1 | 12 | 50,000 | 7.37 | 73.67 |
| Cattle | Muscle | A | 1 | 12 | 100,000 | 4.15 | 41.53 |
| Cattle | Muscle | B | 1 | 12 | 70,000 | 4.08 | 40.84 |
| Cattle | Spleen | A | 1 | 12 | 50,000 | 1.17 | 11.67 |
| Cattle | Spleen | B | 1 | 12 | 50,000 | 9.59 | 95.86 |
| Pig | Adipose | A | 1 | 12 | 500,000 | 3.42 | 34.22 |
| Pig | Adipose | B | 1 | 12 | 500,000 | 4.52 | 45.24 |
| Pig | Cerebellum | A | 1 | 10 | 200,000 | 1.28 | 12.85 |
| Pig | Cerebellum | B | 1 | 10 | 500,000 | 0.91 | 9.09 |
| Pig | Cerebellum | B | 2 | 11 | 200,000 | 0.77 | 7.66 |
| Pig | Cortex | A | 1 | 10 | 200,000 | 1.48 | 14.75 |
| Pig | Cortex | B | 1 | 10 | 200,000 | 1.53 | 15.34 |
| Pig | Hypothalamus | A | 1 | 12 | 500,000 | 0.47 | 4.68 |
| Pig | Hypothalamus | A | 2 | 11 | 500,000 | 0.90 | 8.98 |
| Pig | Hypothalamus | B | 1 | 12 | 500,000 | 1.41 | 14.10 |
| Pig | Hypothalamus | B | 2 | 11 | 500,000 | 0.37 | 3.65 |
| Pig | Liver | A | 1 | 10 | 200,000 | 0.71 | 7.06 |
| Pig | Liver | B | 1 | 10 | 500,000 | 2.11 | 21.08 |
| Pig | Lung | A | 1 | 10 | 200,000 | 2.00 | 20.00 |
| Pig | Lung | B | 1 | 10 | 500,000 | 3.04 | 30.36 |
| Pig | Muscle | A | 1 | 10 | 500,000 | 6.26 | 62.62 |
| Pig | Muscle | B | 1 | 10 | 500,000 | 3.16 | 31.65 |
| Pig | Spleen | A | 1 | 10 | 500,000 | 7.04 | 70.41 |
| Pig | Spleen | B | 1 | 10 | 200,000 | 2.34 | 23.40 |

**Table S2.**
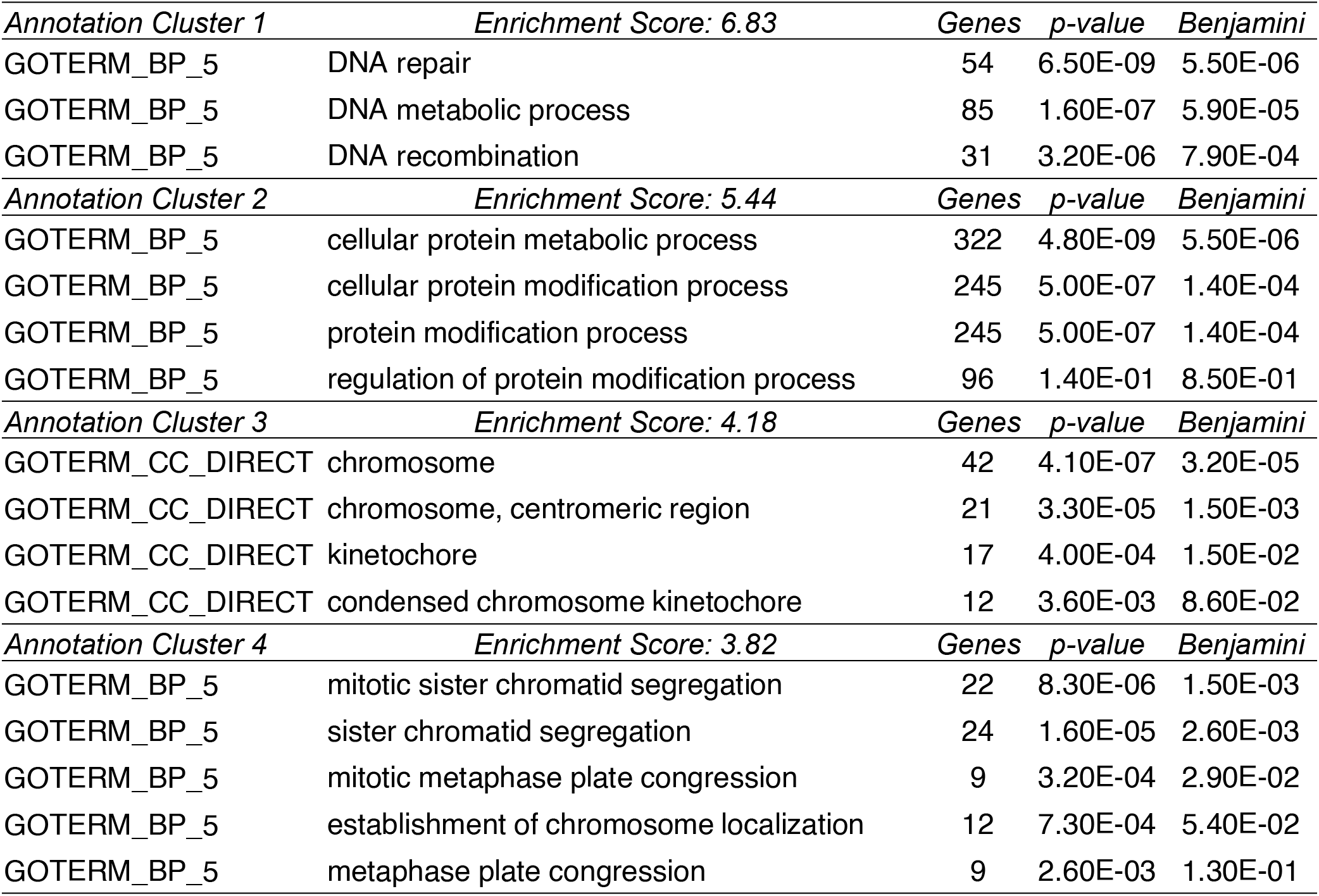
Functional annotation clustering of genes with conserved and global TSS accessibility. Genes with accessible TSS (± 50 bp) in all profiled tissues in all species were subjected to functional annotation clustering to identify enriched cellular functions. Top four clusters reported.

| *Annotation Cluster 1* |  | *Enrichment Score: 6.83* |  | *Genes* | *p-value* | *Benjamini* |
| --- | --- | --- | --- | --- | --- | --- |
| GOTERM_BP_5 | DNA repair |  |  | 54 | 6.50E-09 | 5.50E-06 |
| GOTERM_BP_5 | DNA metabolic process |  |  | 85 | 1.60E-07 | 5.90E-05 |
| GOTERM_BP_5 | DNA recombination |  |  | 31 | 3.20E-06 | 7.90E-04 |
| *Annotation Cluster 2* |  | *Enrichment Score: 5.44* |  | *Genes* | *p-value* | *Benjamini* |
| GOTERM_BP_5 | cellular protein metabolic process |  |  | 322 | 4.80E-09 | 5.50E-06 |
| GOTERM_BP_5 | cellular protein modification process |  |  | 245 | 5.00E-07 | 1.40E-04 |
| GOTERM_BP_5 | protein modification process |  |  | 245 | 5.00E-07 | 1.40E-04 |
| GOTERM_BP_5 | regulation of protein modification process |  |  | 96 | 1.40E-01 | 8.50E-01 |
| *Annotation Cluster 3* |  | *Enrichment Score: 4.18* |  | *Genes* | *p-value* | *Benjamini* |
| GOTERM_CC_DIRECT | chromosome |  |  | 42 | 4.10E-07 | 3.20E-05 |
| GOTERM_CC_DIRECT | chromosome, centromeric region |  |  | 21 | 3.30E-05 | 1.50E-03 |
| GOTERM_CC_DIRECT | kinetochore |  |  | 17 | 4.00E-04 | 1.50E-02 |
| GOTERM_CC_DIRECT | condensed chromosome kinetochore |  |  | 12 | 3.60E-03 | 8.60E-02 |
| *Annotation Cluster 4* |  | *Enrichment Score: 3.82* |  | *Genes* | *p-value* | *Benjamini* |
| GOTERM_BP_5 | mitotic sister chromatid segregation |  |  | 22 | 8.30E-06 | 1.50E-03 |
| GOTERM_BP_5 | sister chromatid segregation |  |  | 24 | 1.60E-05 | 2.60E-03 |
| GOTERM_BP_5 | mitotic metaphase plate congression |  |  | 9 | 3.20E-04 | 2.90E-02 |
| GOTERM_BP_5 | establishment of chromosome localization |  |  | 12 | 7.30E-04 | 5.40E-02 |
| GOTERM_BP_5 | metaphase plate congression |  |  | 9 | 2.60E-03 | 1.30E-01 |

**Table S3.** Functional annotation clustering of genes near conserved intergenic open chromatin. Genes that were closest (within 100kb) to intergenic open chromatin that was conserved in all three species were subjected to functional annotation clustering to identify enriched cellular functions. Top four clusters reported.

| *Annotation Cluster 1* |  | *Enrichment Score: 5.93* | *Genes* | *p-value* | *Benjamini* |
| --- | --- | --- | --- | --- | --- |
| GOTERM_BP_5 | nervous system development |  | 41 | 1.30E-07 | 2.60E-04 |
| GOTERM_BP_5 | central nervous system development |  | 24 | 9.50E-07 | 9.30E-04 |
| GOTERM_BP_5 | brain development |  | 19 | 1.30E-05 | 4.30E-03 |
| *Annotation Cluster 2* |  | *Enrichment Score: 3.48* | *Genes* | *p-value* | *Benjamini* |
| GOTERM_BP_5 | mesenchymal cell development |  | 9 | 5.20E-05 | 1.10E-02 |
| GOTERM_BP_5 | mesenchymal cell differentiation |  | 9 | 8.70E-05 | 1.20E-02 |
| GOTERM_BP_5 | mesenchyme development |  | 10 | 1.10E-04 | 1.40E-02 |
| GOTERM_BP_5 | ameboidal-type cell migration |  | 11 | 3.00E-04 | 2.50E-02 |
| GOTERM_BP_5 | neural crest cell migration |  | 5 | 9.10E-04 | 4.80E-02 |
| GOTERM_BP_5 | central nervous system neuron differentiation |  | 8 | 1.10E-03 | 5.40E-02 |
| GOTERM_BP_5 | neural crest cell development |  | 5 | 2.90E-03 | 1.10E-01 |
| *Annotation Cluster 3* |  | *Enrichment Score: 3.26* | *Genes* | *p-value* | *Benjamini* |
| GOTERM_BP_5 | regionalization |  | 12 | 7.70E-05 | 1.10E-02 |
| GOTERM_BP_5 | embryo development ending in birth or egg hatching |  | 17 | 1.50E-04 | 1.50E-02 |
| GOTERM_BP_5 | neural tube development |  | 6 | 1.50E-02 | 2.60E-01 |
| *Annotation Cluster 4* |  | *Enrichment Score: 2.78* | *Genes* | *p-value* | *Benjamini* |
| GOTERM_BP_5 | organ morphogenesis |  | 24 | 3.10E-06 | 1.50E-03 |
| GOTERM_BP_5 | cell migration |  | 22 | 2.60E-04 | 2.30E-02 |
| GOTERM_BP_5 | ameboidal-type cell migration |  | 11 | 3.00E-04 | 2.50E-02 |
| GOTERM_BP_5 | cell development |  | 33 | 3.30E-04 | 2.50E-02 |
| GOTERM_BP_5 | circulatory system development |  | 20 | 4.10E-04 | 2.70E-02 |
| GOTERM_BP_5 | cardiovascular system development |  | 20 | 4.10E-04 | 2.70E-02 |
| GOTERM_BP_5 | heart development |  | 14 | 6.70E-04 | 3.80E-02 |
| GOTERM_BP_5 | vasculature development |  | 13 | 5.40E-03 | 1.60E-01 |
| GOTERM_BP_5 | blood vessel development |  | 12 | 9.50E-03 | 2.10E-01 |
| GOTERM_BP_5 | positive regulation of cell migration |  | 9 | 2.00E-02 | 3.10E-01 |
| GOTERM_BP_5 | positive regulation of cell motility |  | 9 | 2.40E-02 | 3.20E-01 |
| GOTERM_BP_5 | positive regulation of cellular component movement |  | 9 | 2.70E-02 | 3.40E-01 |
| GOTERM_BP_5 | angiogenesis |  | 7 | 1.20E-01 | 6.50E-01 |

**Table S4.**
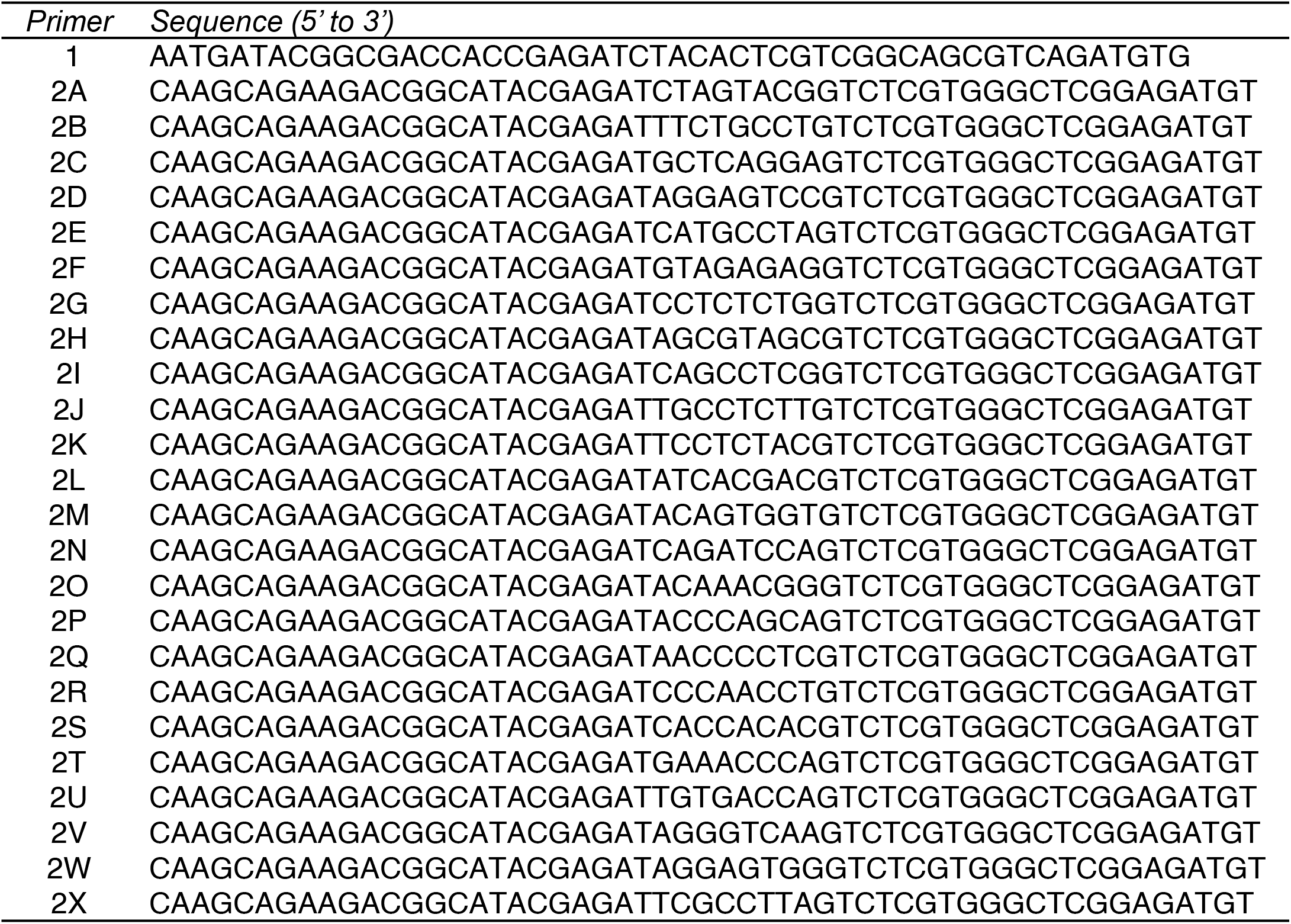
ATAC-seq oligos used for PCR. Sequences have been previously described by Buenrostro *et al*, 2013. Primers 2A-2X contain variable barcodes which permit library pooling prior to sequencing, and which were used to demultiplex sequencing data.

